# Arabidopsis produces distinct subpopulations of extracellular vesicles that respond differentially to biotic stress

**DOI:** 10.1101/2024.04.18.589804

**Authors:** Benjamin L. Koch, Brian D. Rutter, Roger W. Innes

## Abstract

Extracellular vesicles (EVs) secreted by mammalian cells are highly heterogenous in contents and function. Whether this is also true for EVs secreted by plant cells is not yet known. To address this knowledge gap, we used high-resolution density gradient ultracentrifugation to separate distinct subpopulations of Arabidopsis EVs. We analyzed the protein content, morphology, and purity of these subpopulations, confirming the presence of three distinct EV subpopulations. The EV marker protein TETRASPANIN 8 (TET8) was detected only in medium-density EVs and was not associated with cell wall nanofilaments, which was unique among EV proteins. TET8 and PENETRATION 1 (PEN1) were confirmed to be secreted on mostly separate EV populations using total internal fluorescence microscopy. We found that EV marker proteins are differentially secreted in response to phytohormones, changes in growth temperature, and infection with fungal pathogens *Colletotrichum* and *Golovinomyces cichoracearum*. EV subpopulations marked by TET8, PEN1, and RPM1-INTERACTING PROTEIN 4 (RIN4) were highly increased as soon as one day after fungal infection, while other EV populations remained unaffected. Together these data indicate that Arabidopsis EVs are highly heterogenous and suggest that specific EV subpopulations may contribute to plant immunity.

## Introduction

Extracellular vesicles (EVs) are produced by all kingdoms of life (Gill et al., 2019). A key feature of EVs, particularly in human cell culture, is their diversity (Mathieu et al., 2019). EVs range in diameter from 50 nm to 5 µm, and variations in size can be partially attributed to differences in biogenesis (Jeppesen et al., 2023). EVs are produced and secreted by at least three different pathways: fusion of a multivesicular body with the plasma membrane, budding of microvesicles directly from the plasma membrane, and production of large apoptotic bodies during cell death. EV marker proteins, as well as lipid composition and RNA content, help distinguish different EV populations from one another. Distinct protein, lipid, and RNA content may enable specific populations of EVs to perform different physiological functions. For example, in EVs derived from tumors, enrichment of specific EV transmembrane proteins can determine which organs are targeted by metastasizing tumors (Hoshino et al., 2015). Similarly, fibroblast cells become more invasive when treated with EVs isolated from colon carcinoma cells, and small exosomes cause a greater increase in invasiveness than larger microvesicles (Xu et al., 2015). As a third example, the secretion of cytokines from T lymphocytes can be differentially modulated by treatment with small and large EVs from immature dendritic cells (Tkach et al., 2017).

Recent studies have isolated and characterized EVs of Arabidopsis and other plant species. There is growing evidence that plant EVs have evolved to play a role in plant immunity (Stotz et al., 2022; Rutter and Innes, 2018; Rybak and Robatzek, 2019). Plant EV secretion is increased during bacterial infection, and upon stimulation with the immune signaling hormone salicylic acid (SA) (Rutter and Innes, 2017). EV secretion is increased in response to the bacterial pathogen *Pseudomonas syringae* and the fungal necrotrophic pathogen *Botrytis cinerea* (He et al., 2021; Rutter and Innes, 2017). It has also been posited that plant EVs contain RNA that is delivered to pathogens, but this topic is still debated (Baldrich et al., 2019; Cai et al., 2018; Zand Karimi et al., 2022). Plant EVs are enriched in defense-related proteins (Rutter and Innes, 2017). A recent report suggested that the EV proteins PENETRATION 1 (PEN1) and TETRASPANIN 8 (TET8) label distinct populations of plant EVs, and that these populations could be separated by differential ultracentrifugation (He et al., 2021).

We set out to test the hypothesis that there are different populations of plant EVs and that these distinct EV populations have unique functions. After isolating EVs from the apoplast of Arabidopsis plants using differential ultracentrifugation, pelleted EVs were fractionated using a high-resolution density gradient. Probing the density gradient fractions by immunoblot revealed three populations of EVs characterized by a distinct pattern of EV marker proteins. We characterized the response of these EV populations to biotic and abiotic stress by modulating immune hormones salicylic acid and jasmonic acid, changing growth temperatures, and by infecting with the biotrophic powdery mildew fungus *Golovinomyces cichoracearum* and hemibiotrophic fungus *Colletotrichum higginsianum*. We conclude that there are distinct plant EV populations, and that secretion of these populations are differentially regulated in response to biotic and abiotic stress. We speculate that different populations of EVs are created either through different biogenesis pathways, or that EV proteins are differentially loaded during stress conditions such as fungal infection.

## MATERIALS AND METHODS

### Plant materials

The Arabidopsis genotypes used in this study were: Columbia-0; *sid2Δ*; *coi1-16*; 35S::RFP-PEN1 crossed with TET8promoter::TET8-GFP; HA-AGO2; and *pad4*. Arabidopsis plants were grown in a growth room or growth chamber at 22°C, 60% relative humidity, and under 150 µmol m^-2^ s^-1^ light for 9 hours per day. Arabidopsis seeds were sterilized using 70% v/v ethanol then 20% v/v bleach for 5 min, washed with sterile water 5 times, vernalized for at least 2 days at 4°C in darkness, and germinated on ½ strength MS media 1% agar pH 5.8 (Research Products International, #M10200) in the same growing conditions as Arabidopsis plants. 10- to 14-day old seedlings were transferred to soil (Sungro Sunshine Mix #5) and grown beneath a humidity dome for 7 days before removing the dome. Plants were watered with 1 liter of water plus fertilizer (Miracle Gro, all-purpose plant food, water-soluble) when the soil appeared dry.

### Isolation of apoplastic wash fluid

Apoplastic wash fluid (AWF) was isolated using a previously published protocol (Rutter et al., 2017; Welsh et al., 2024). In brief, 6-8-week-old Arabidopsis rosettes were harvested by cutting off the root. For all experiments, except fungi infection, rosettes were washed in distilled water three times. Rosettes were then infiltrated by placing in a beaker abaxial side facing up, submerging in 4°C vesicle isolation buffer (VIB, 20 mM MES, 2 mM CaCl_2_, and 10 mM NaCl, pH 6) using a French press coffee maker, placing submerged plants in a vacuum chamber, and applying a vacuum using a vacuum pump (Thermo Savant VP 100 Two Stage Model #1102180403) for 30 seconds until 95% of leaf area appeared infiltrated. Infiltrated rosettes were patted dry gently with paper towel to remove all excess buffer. 2-3 rosettes were gently placed inside modified 50 mL syringes with 10-20 extra holes melted into the bottom to allow buffer to freely flow through. The loaded syringes were placed in 250 mL wide-mouth bottles, using parafilm wrapped around the syringe as an adaptor where necessary to hold the syringe in the bottle mouth. Bottles were placed in a fixed-angle rotor (Beckman JA-14 #339247) and centrifuged at 700 g for 30 min at 4°C using slow acceleration to isolate AWF from the leaves. AWF was passed through a 0.22 µm filter (PALL Life Sciences #4612, EMD Millipore #SCGP00525).

### EV pellet isolation

Filtered AWF was centrifuged in a fixed-angle rotor (Beckman TLA100.3 [#349490], TLA100.4, or TLA110 [#366735]) using a benchtop ultracentrifuge (Beckman Optima TLX-120) in polycarbonate tubes (Beckman 13 x 51 mm #349622) at 10,000 g for 30 min at 4°C. The supernatant was transferred to a new tube by pipetting. For differential ultracentrifugation (Figure 1), the supernatant was centrifuged at 40,000 g for 1 h at 4°C. The supernatant was moved to a new tube by decanting, all residual supernatant was removed by wicking off with a Kimwipe, and the pellet (P40) was resuspended in 20 mM Tris HCl pH=7.5 prepared by diluting Tris stock solution in Millipore water and filtered through a 0.22 micron filter immediately prior to use (ultraclean Tris resuspension buffer). The P40 supernatant was centrifuged at 100,000 g for 1hr at 4°C. The supernatant was removed by decanting, and the pellet (P100 minus P40, P100-40) was resuspended as before. For experiments where a P100 was used, the P10 supernatant was spun at 100,000 g for 1 h at 4°C and resuspended as before – no 40,000 g spin is used when “P100” is indicated.

**FIGURE 1.**
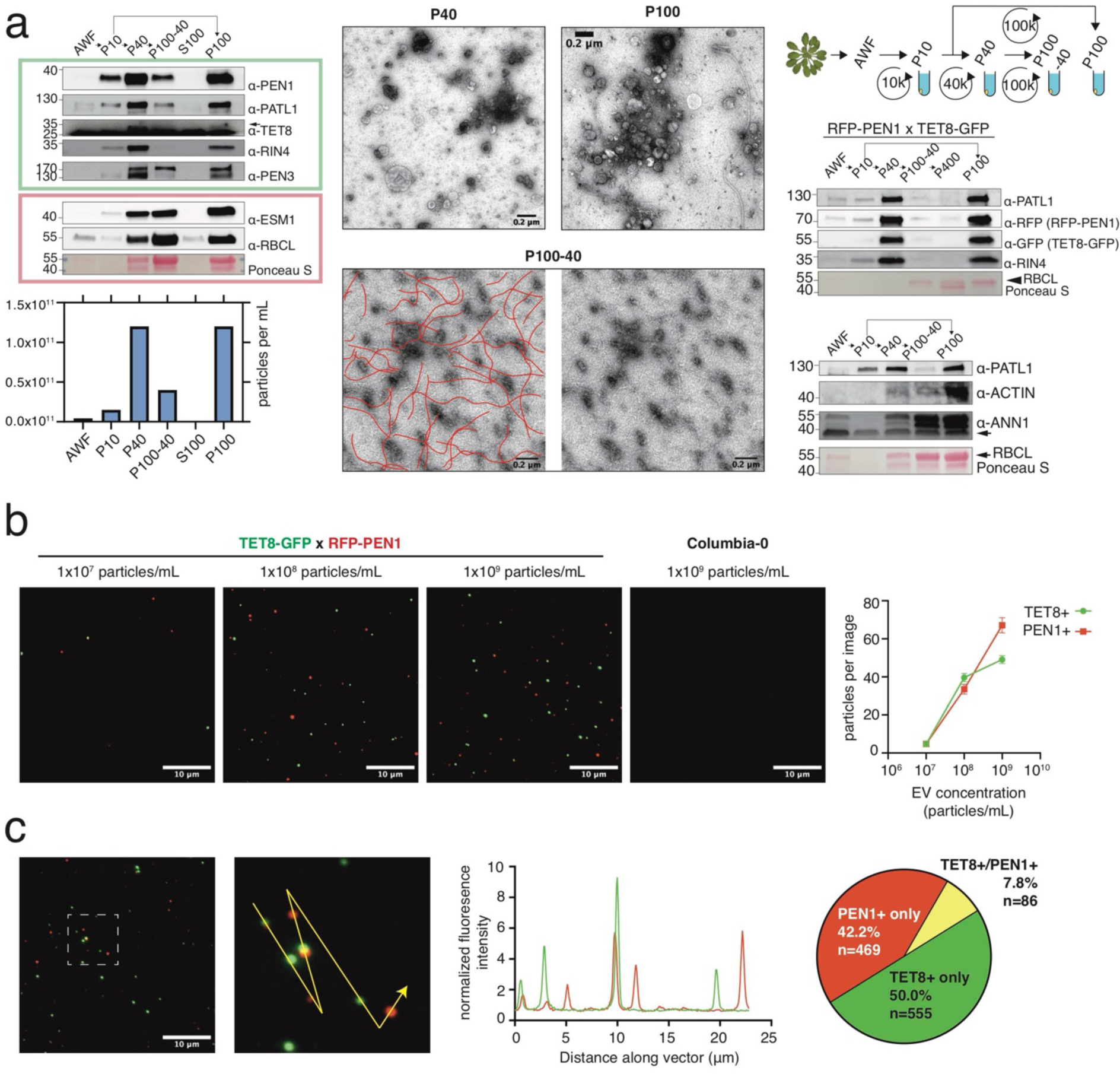
TET8 and PEN1 mark largely dissimilar EV populations. (a) Immunoblots, nanoparticle tracking analysis (NTA), and representative transmission electron microscopy (TEM) images of different extracellular extracts from Arabidopsis rosettes processed by differential ultracentrifugation, as indicated in diagram. For immunoblot, samples are apoplastic wash fluid (AWF), pellets collected from 5 mL equivalent volume of AWF. Numbers on left of immunoblots indicate the approximate kiloDalton (kDa). Since the TET8 antibody PHY1490A strongly cross-reacts with a <25kDa protein abundant in EV pellets, the arrowhead indicates the TET8-specific band that corresponds to the predicted molecular weight of the protein and verified by probing a *tet8* null mutant (Figure S1). For TEM, red lines highlight nanofilaments abundant in the P100-P40 image. Arabidopsis expressing RFP-PEN1 and TET8-GFP were used where indicated. P10 = pellet from 10,000 g centrifugation of AWF; P40 = pellet from 40,000 g centrifugation of P10 supernatant; P100-40 = pellet from 100,000 g centrifugation of P40 supernatant; P100 = pellet from 100,000 g centrifugation from P10 supernatant (no 40,000 g step); P400 = pellet from 400,000 g centrifugation of P100 supernatant; S100 = supernatant after 100,000 g spin. (b,c) Total internal reflection fluorescence microscopy (TIRF-M) of P40 EVs. Representative merged images with TET8-GFP colored green and RFP-PEN1 colored red. Particle concentration was determined by NTA. The number of TET8+ and PEN1+ EVs per field of view and the number of co-localized particles was determined using the object-based co-localization Fiji plugin DiAna (Gilles et al., 2017). Objects closer than 200 nm were defined as co-localized. Error bars represent ± standard error of the mean (SEM). Size of selected area for line intensity scan: 10 µm x 10 µm. In pie chart, n indicates the number of particles which belong to each category (1,110 total particles were analyzed).

**FIGURE 2.**
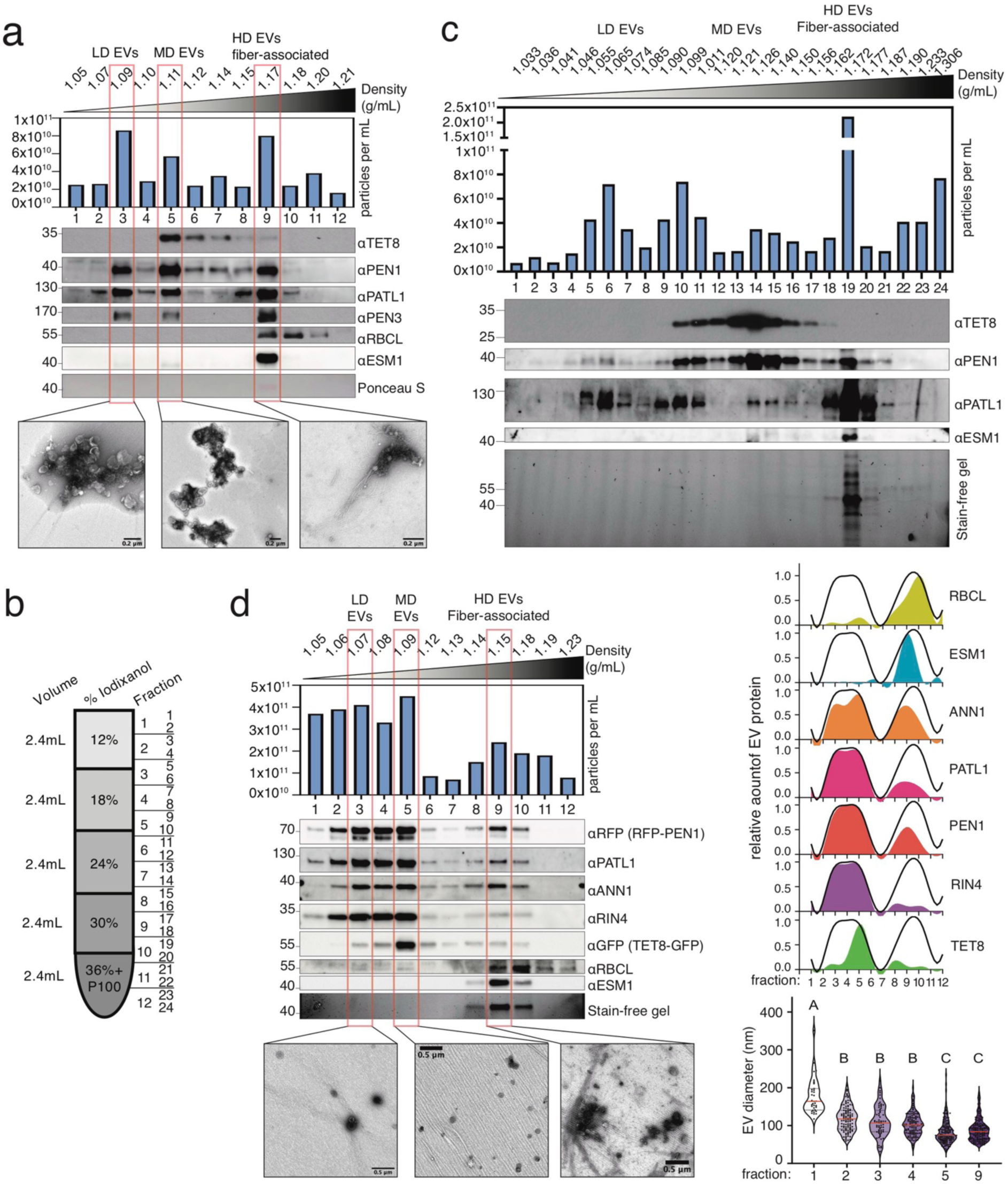
High-resolution density gradient separates distinct subpopulations of Arabidopsis EVs. Immunoblots, NTA, and representative TEM images of 12 (a,d) or 24 (c) fractions of a discontinuous iodixanol gradient layered above a P100 EV pellet shown in (b). The density of each fraction was determined using refractometry, or by measuring mass and volume. Relative amount of EV protein (d) was determined using densitometry, and the black line shows the maximum signal at any given fraction. EV diameter (d) was determined by measuring the diameter of each particle with EV morphology three times and calculating the mean diameter for each particle. Number of EVs counted for fraction 1=34, fraction 2=105, fraction 3=64, fraction 4=120, fraction 5=140, and fraction 9=159. The red line indicates the average diameter, and the dotted black lines indicate the interquartile range. Average EV diameter of fraction1=178nm; fraction 2=118nm; fraction 3=110nm; fraction 4=103nm; fraction 5=83nm; fraction 9=88nm. Letters indicate statistical significance determined by two-way ANOVA with post-hoc Tukey multiple comparison. LD EVs=low-density EVs, MD EVs=medium-density EVs, HD EVs=high-density EVs.

### Total internal reflection fluorescence microscopy (TIRF-M)

P40 pellets were prepared as above by infiltrating transgenic and wild type Arabidopsis with VIB supplemented with 10 mM EGTA to prevent clumping of particles, followed by filtration and differential ultracentrifugation of AWF, and resuspension of P40 in ultraclean Tris resuspension buffer. P40 pellets were not frozen between isolation and microscopic observation. Slides and coverslips (22 x 22 mm, No. 1.5) were cleaned with 100% ethanol and dried for several hours in a 60°C oven. 10 µL of sample was pipetted onto the surface of the slide beneath a clean coverslip, sealed with nail polish, and observed with a GE Deltavision OMX SR super-resolution microscope using the Ring TIRF Option with 1.1516 refractive index immersion oil. After the focal plane was found for each mounted sample, a new area of the same sample was imaged to avoid photobleaching. The same laser excitation, image acquisition settings, and Fiji image processing were used for transgenic and wild type EVs. The Fiji plugin DiAna was used for object-based co-localization analyses and counting the number of particles per field of view (Gilles et al., 2017). The center of fluorescent objects was determined with Gaussian methods and particles whose centers were closer than 200nm were considered co-localized. Particle concentration for dilution series was determined by nanoparticle tracking analysis (NTA). Statistical analyses were performed in Excel (Microsoft) and Graphpad Prism. For phalloidin staining, rhodamine-phalloidin (Invitrogen #R415) was resuspended in DMSO at a concentration of 66 µM and mixed with the EV pellet to a final concentration of 3.3 µM 5 minutes before mounting the sample on a slide and imaging as above.

### Density gradient

Filtered AWF was distributed into tubes (Beckman Ultra-Clear 14 x 89 mm, #344059 or 25 x 89 mm, #344058) and centrifuged in a swinging bucket rotor (Beckman SW 41 Ti #331362 or SW 32 Ti #369650, respectively) in an ultracentrifuge (Beckman Optima XPN-100 #A99846) at 10,000 g for 30 min at 4°C. The supernatant was moved to a new tube by pipetting, and centrifuged at 100,000 g for 1 h at 4°C to pellet EVs (P100). The supernatant was removed by pipetting to avoid disturbing the pellet, and the pellet was resuspended in VIB. The density gradient layers were made by mixing Optiprep (60% iodixanol) with VIB, except for the bottom layer which was made my mixing Optiprep with the resuspended P100. The density gradient was made by transferring the layers sequentially into a tube (Beckman Ultra-Clear 14 x 89 mm, #344059) using a transfer pipet being careful to avoid mixing the layers. The density gradient was centrifuged at 100,000 g for 17 h at 4°C in an SW 41 Ti rotor. Fractions (1 mL or 0.5 mL, for low-/high-resolution and 24-part gradients respectively) were carefully removed starting from the top of the gradient using a pipet and transferred to a polycarbonate tube (Beckman 13 x 51 mm, #349622). The density of each fraction was determined either by weighing the fraction and dividing by the volume, or by refractometry (Misco Palm Abbe #PA201x). Fractions were diluted to 3 mL using VIB, mixed with a pipet, and centrifuged at 100,000 g for 2 h at 4°C. The supernatant was removed by decanting, and the pellet was resuspended in ultraclean Tris resuspension buffer.

### Enzymatic digestion of EV pellets

For protease protection assays (Figure 3), EV pellets were resuspended in ultraclean 150 mM Tris HCl pH=7.8 buffer and aliquoted. EV pellets were not frozen between isolation and performing protease protection assay. For detergent treatment, 6% v/v Triton X-100 (EMD Millipore #TX1568-1) added by pipetting for a final concentration of 1% TX-100, and the sample was incubated for 30 min at 4°C. For protease treatment, 10 µg/mL stocks of trypsin (Promega #V5113) and/or chymotrypsin (Sigma-Aldrich #C4129) were added to final concentrations of 1-2 µg/mL, and the sample was incubated for 1 h or 2 h at 37°C, as indicated in Figure 3.

**FIGURE 3.**
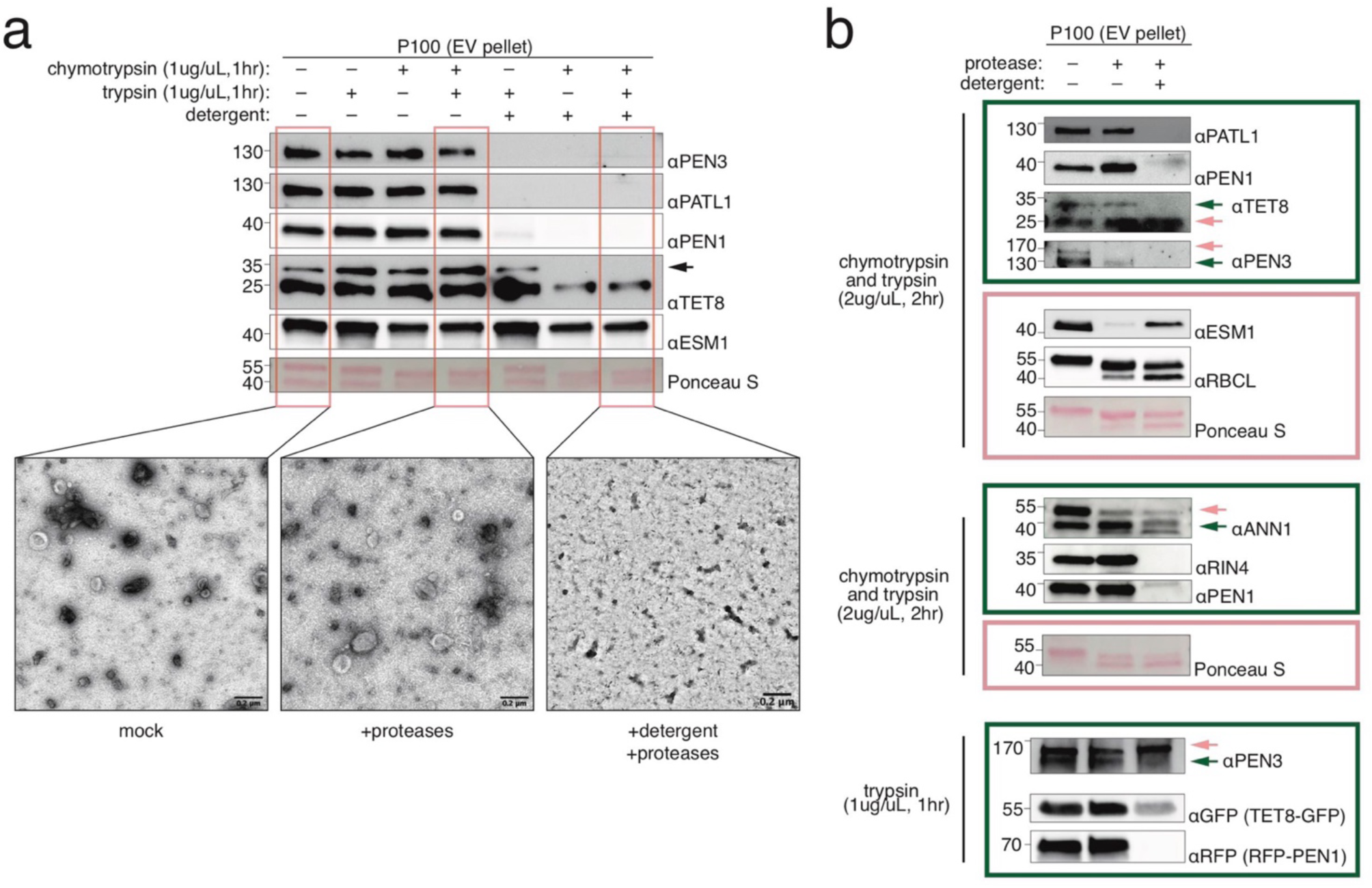
EV proteins are protected from protease digestion. Immunoblot and representative TEM images of P100 EV pellets with protease and detergent treatments. Detergent treatments were 1% Triton-X100 mixed with EVs by pipetting and incubated at 4°C for 30 min. Protease treatments were performed after detergent treatments. Protease treatments were at 37°C, pH=7.8 using varying concentration and times indicated in the figure. For (b), Dark green boxes and dark green arrows indicate EV proteins that were made susceptible to protease digestion only after pre-treatment with detergent. Light red boxes and light red arrows indicate non-EV proteins that were susceptible or partially susceptible to protease digestion regardless of detergent treatment. These immunoblots also serve to validate EV antibodies.

For digestion of nanofilaments (Figure 4), EV pellets were resuspended in VIB with 10 mM EGTA adjusted to pH=5 and passed through a 0.22 µm filter immediately prior to use. Cellulase Onozuka R-10 (Dot Scientific #DSC32200-5) and Pectolyase Y-23 (MP Biomedicals #320951) were resuspended in the same buffer at 10% w/v and 1% w/v respectively. EV pellets were digested with these enzymes at 37°C at the concentrations and times indicated in Figure 4.

**FIGURE 4.**
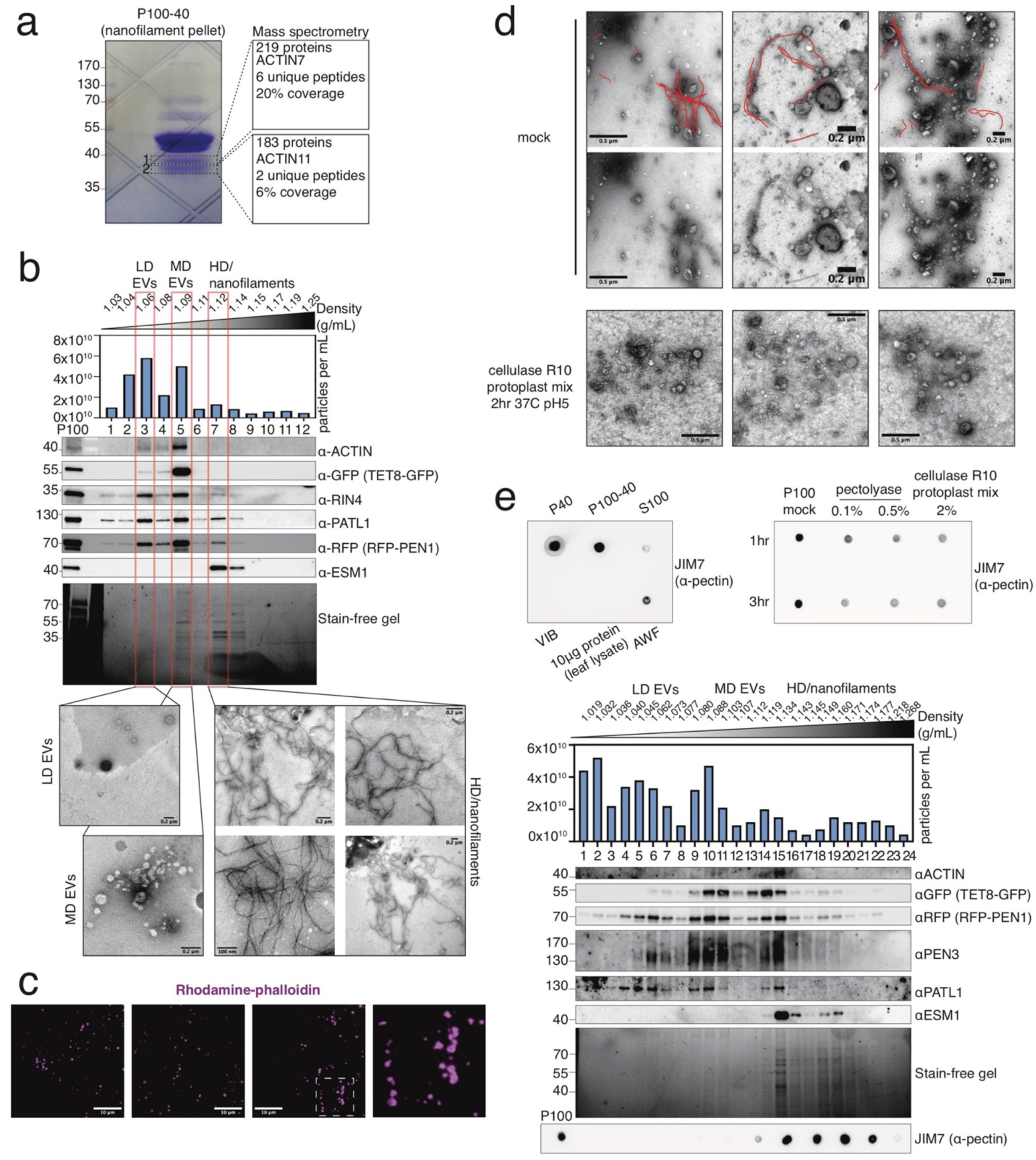
Nanofilaments derived from the cell wall co-pellet with Arabidopsis EVs. (a) Coomassie-stained SDS-PAGE of P100-40 pellet enriched in nanofilaments. Gel fragments labeled 1 and 2 corresponding to abundant proteins around 40 kDa that were excised and processed using mass spectrometry. (b) Immunoblot, NTA, and representative TEM images of high-resolution density gradient separation of EVs isolated in VIB containing 10 mM EGTA. Addition of EGTA caused HD EVs to equilibrate in fraction 7 instead of fraction 9, as well as helping with observation of nanofilaments. Density of fractions was determined using refractometry. (c) TIRF-M of P40 EVs stained with rhodamine-phalloidin, a filamentous actin dye. Three representative images are shown, and one selected area is enlarged, size of selected area is 10 µm x 10 µm. (d) Representative TEM images of P100 digested with cellulase R10 Onozuka protoplasting enzyme mix (cellulase: 1.0 U/mg, pectinase 0.4 U/mg, hemicellulase 1.0 U/mg, alpha-Amylase 0.6 U/mg, protease: 0.01 U/mg) at indicated conditions or mock treatment. Red lines highlight nanofilaments derived from the cell wall, since the filaments are not present after digestion with the cell wall enzyme mix. (e) Dot blot detection of pectin in EV pellets and in a high-resolution density gradient. The pectin antibody JIM7 detects equivalent amounts of pectin in P40 and P100-40, consistent with the observation of cell wall nanofilaments in both these EV pellets (see Figure 1). P40 and P100-40 were collected from 2.5 mL AWF. Digestion of the P100 EV pellet with pectolyase or cell wall enzyme mix confirms that JIM7 recognizes pectin. The P100 for each treatment was collected from the equivalent of 0.5mL AWF. For high-resolution density gradient separation of EVs, 10 mM EGTA was used in the VIB buffer (similar to b). In contrast to most EV markers, which equilibrate at low- and medium-densities, pectin equilibrates at high-densities, consistent with the observation of nanofilaments at high-densities.

### Nanoparticle Tracking Analysis (NTA) and Transmission Electron Microscopy (TEM)

Resuspended EV pellets, digested EV pellets, or density gradient pellets were not frozen prior to observation using NTA or TEM. For NTA, the resuspended pellets were diluted in ultraclean Tris resuspension buffer. The nanoparticle tracker (Particle Metrix PMX 120 Zetaview Mono 488 Laser running Zetaview software version 8.05.16 SP3) was calibrated with 100 nm beads (nanoComposix silica nanospheres #SISN100). Diluted EV samples (ranging from 1:500 to 1:2000 dilutions) were injected such that approximately 100 particles were visible per field of view could be observed when using the camera settings of 75 sensitivity, 30 frame rate, and 100 shutter. Particle concentration was recorded as the average of 11 camera positions. For TEM, copper or nickel grids formvar-coated grids (Electron Microscopy Sciences, # FCF300) were prepared using glow discharge at 0.4 mBar and 15 mA for 1 min (PELCO easiGlow #91000 and #91040). 5 µL of sample (diluted 1:5 where necessary) was pipetted onto the formvar-coated surface and allowed to adhere for 5 minutes. Sample was wicked from the grid using Whatman filter paper and stained twice with 10 µL 2% uranyl acetate by pipetting the uranyl acetate across the surface of the grid. Stained grids were protected from light and allowed to dry at least 16 h before imaging at 80kV using a JEOL JEM-1010 transmission electron microscope.

### Protein preparation and Immunoblot

After AWF extraction, leaves were flash frozen in liquid nitrogen and ground into a fine powder. Cold protein extraction buffer (150 mM NaCl, 50 mM Tris HCl pH 7.5, 0.1% v/v Nonidet-P40, 1% v/v plant protease inhibitor cocktail) was added and mixed with sample on ice using a mechanical pestle for 1 minute. After incubation on ice for 20 minutes, debris was pelleted twice by centrifuging at 16,000 g for 10 minutes at 4°C. Protein concentrations of the cell lysate or EV pellets were quantified with Pierce 660 nm reagent (Sigma #22660) using the manufacturer’s protocol and comparing to a bovine serum albumin standard curve.

For SDS-PAGE, equal mass of cell lysate protein diluted in protein extraction buffer and EV pellets isolated from equivalent volumes of AWF were mixed with 5X loading dye (250 mM Tris HCl pH 6.8, 10% w/v SDS, 40% v/v glycerol, 0.02% bromoethanol blue, 5% v/v 2-mercaptoethanol) and incubated at 95°C for 5 minutes. Samples and protein ladder (PageRuler Prestained Protein Ladder, Thermo #26616) were run in pre-cast 4– 20% TGX stain-free denaturing protein gels (Biorad #4568093, #4568096, #5678094, #5678095) in a minigel system or criterion system (Biorad #1658005, #1656001) using running buffer (25 mM Tris, 200 mM glycine, 3.5 mM SDS) for 30 min at 50V followed by 1 h at 120V, or until samples had migrated to the end of the gel. Stain-free gel images were captured in Chemidoc MP imaging system (Biorad #12003154) after 45 seconds of activation with UV light. Protein was transferred to 0.45 µm nitrocellulose membrane (Cytiva Amersham Protran #10600003) using a tank system (Biorad #1703930, #1704070) in transfer buffer (25 mM Tris, 200 mM glycine, 20% v/v methanol) for 50 minutes at 300 mA. Protein transfer to the membrane was visualized by applying Ponceau Stain (0.1% w/v Ponceau S Stain, 5% v/v acetic acid) for 5 minutes and rinsing in distilled water.

For immunoblot, membranes were blocked in 5% skim milk dissolved in Tris-buffered saline + Tween 20 (TBST, 100 mM Tris, 150 mM NaCl, 0.1% Tween-20, pH 7.5) for 1 h at 25°C with gentle rocking. After blocking, membranes were incubated with primary antibody (see Table 2) for 16 h at 4°C. After primary antibody, membranes were washed with TBST 3 times for 10 minutes each, then secondary antibody was applied where applicable for 1-2 h at 25°C and washed with TBST 3-5 times for 10 min each. Enhanced chemiluminescent substrate (National Diagnostics Protoglow #CL-300) was applied to membranes for 5 min, and signal was detected using a Chemidoc MP imaging system (Biorad #12003154) or by exposing membrane to film, developed using a table-top x-ray film processer (Konica #SRX101A), and scanned at 600dpi. Immunoblot images were formatted in Adobe Illustrator. Densitometry of immunoblot images was calculated using FIJI. Graphs and statistical analyses were produced in Graphpad Prism. The specificity of antibodies was confirmed by probing Arabidopsis null mutants for loss of the appropriate band (Figure S1).

For dot blotting, 5 µL droplets of sample were pipetted onto the surface of 0.45 µm nitrocellulose membrane (Cytiva Amersham Protran #10600003). For some samples, as many as three 5 µL droplets were applied to the membrane in succession. Membranes were blocked in 5% w/v skim milk dissolved in phosphate-buffered saline + Tween 20 (PBST, 10 mM Na_2_HPO_4_, 1.8 mM KH_2_PO_4_, 137 mM NaCl, 2.7 mM KCl, 0.02% Tween-20, pH 7.4) for 1 h at 25°C with gentle rocking. After blocking, membranes were incubated with JIM7 pectin antibody (see Table 2) for 16 h at 4°C. After primary antibody, membranes were washed with PBST 3 times for 10 min each, then secondary antibody was applied for 1 h at 25°C and washed with PBST 3 times for 10 min each. Enhanced chemiluminescent substrate (National Diagnostics Protoglow #CL-300) was applied to membranes for 5 minutes, and signal was detected using the Chemidoc MP imaging system (Biorad #12003154).

### Hormone treatment

The mock spray was composed of buffer only (20 mM MES, 0.01% v/v Silwet, pH 6). For salicylic acid, salicylic acid (Sigma #S7401) was dissolved to a concentration of 2 mM in mock buffer. After 1 h of stirring at 25°C to dissolve, pH was reset to 6 using NaOH. For methyl-jasmonic acid, a 20x stock of methyl-jasmonic acid was made by mixing 4.5 µL of methyl-jasmonate (Sigma #392707) with 10 mL of distilled water and rotating for 16 h at 4°C to dissolve. The 20x stock was diluted to 100 µM in mock buffer and stirred for 5 min at 25°C to mix. In a fume hood, 18 plants per treatment were sprayed with 100 mL of mock or hormone solution and watered with an additional 100 mL of solution and covered with a humidity bag to prevent volatile hormones from escaping.

### Fungi and plant infection

The following species were used in this study: *Colletotrichum higginsianum* (*Ch*) isolate IMI349063A; *Colleotrichum destructivum* (*Cd*) isolate LARS 709; and powdery mildew *Golovinomyces cichoracearum* (*Gc*) strain USCS1 (Adam et al., 1999). *Ch* and *Cd* spores were stored at -80°C in 15% v/v glycerol. *Ch* and *Cd* were grown from spore stock in 250 mL Erlenmeyer flasks capped with a foam plug on 100 mL of Mathur’s media (15 mM glucose, 0.15% w/v peptone, 0.05% w/v yeast extract, 5 mM magnesium sulfate heptahydrate, 20 mM potassium dihydrogen phosphate, pH 5.5 with KOH, 2% w/v agar) or ½ PDA (1.2% w/v potato dextrose, pH 5.5 with KOH, 2% w/v agar), respectively, under same conditions as Arabidopsis (see above). After 7-10 days of growth, spores were harvested in distilled water by gently scrubbing the surface of culture vessels with a glass hook. Spores were washed by centrifuging at 500 g for 5 min at 25°C two times. Spores were resuspended in 1 mL distilled water and concentration was determined using a hemocytometer. Spores were diluted to a concentration of 1x10^7^ spores/mL in water. For inoculation, 72 6-week-old Arabidopsis plants were sprayed using an oil mister with 30 mL of spore solution or distilled water as a mock treatment. Sprayed plants were covered with a humidity dome, placed inside a humidity bag, and were replaced in a growth room or chamber under normal growth conditions. *Gc* was maintained on *pad4* plants. For *Gc* inoculation, a settling tower was used. Previously infected leaves from ten *pad4* mutant plants were gently rubbed across a 50 µm nylon mesh placed 46 cm above 108 plants. The inoculum concentration was determined using a hemocytometer at plant level as previously described (Wu et al., 2021), and was found to be approximately 500 spores/mm^2^. The AWF of at least 18 plants was harvested at each time-point.

### Trypan blue staining

Three representative leaves per treatment were placed in trypan blue solution (0.1% w/v trypan blue, 25% v/v lactic acid, 25% v/v phenol [TE buffer equilibrated pH 7.5], 25% v/v glycerol) in 2 mL microfuge tubes, caps were sealed, and tubes were boiled for 1 min using a water bath. Trypan blue solution was removed and replaced with 95% ethanol 3 times over the course of 48 h to destain. After the third destaining step, ethanol was replaced with 2.5 g/mL chloral hydrate solution for at least 16 h. Leaves were mounted on slides in 50% v/v glycerol and mounted leaves were observed using a microscope (Invitrogen EVOS XL Core #AMEX1200).

### Mass spectrometry (MS)

Two independent biological replicates of pelleted low-resolution density gradient fractions were pooled. For protein digestion, samples containing pelleted density gradient fractions were denatured in 8 M urea in 100 mM ammonium bicarbonate. Samples were incubated for 45 min at 57°C with 10 mM Tris(2-carboxyethyl)phosphine hydrochloride to reduce cysteine residue side chains. These side chains were then alkylated with 20 mM iodoacetamide for one hour in the dark at 21°C. The urea was diluted to 1 M urea using 100 mM ammonium bicarbonate. A total of 0.4 µg trypsin (Promega) was added, and the samples were digested for 14 h at 37°C. For MS, the resulting peptide solution was desalted using ZipTip pipette tips (EMD Millipore), dried down and resuspended in 0.1% formic acid. Peptides were analyzed by LC-MS on an Orbitrap Fusion Lumos equipped with an Easy NanoLC1200. Buffer A was 0.1% formic acid in water. Buffer B was 0.1% formic acid in 80% acetonitrile. Peptides were separated on a 90-minute gradient from 0% B to 35% B. Peptides were fragmented by HCD at a relative collision energy of 32%. Precursor ions were measured in the Orbitrap with a resolution of 60,000. Fragment ions were measured in the Orbitrap with a resolution of 15,000. Data were analyzed using Proteome Discoverer (2.5) to interpret and quantify the relative amounts in a label free quantification manner. Data was searched against the Arabidopsis TAIR10 proteome downloaded on 2/12/2015. Trypsin was set as the protease with up to two missed cleavages allowed. Carbamidomethylation of cysteine residues was set as a fixed modification. Oxidation of methionine and protein N-terminal acetylation were set as variable modifications. A precursor mass tolerance of 10 ppm and a fragment ion quantification tolerance of 0.04 Da were used. Data was quantified using the Minora feature detector node within Proteome Discoverer. Proteins with either less than 1 peptide or less than 4% sequence coverage were eliminated from analysis. Protein intensity values were normalized by quantile normalization. To find enrichment of proteins in LD/MD versus HD, the normalized intensity values from LD/MD fraction were divided by the normalized intensity of HD fractions from mock and infected samples respectively. The enrichment values of mock and infected samples were averaged, and the average enrichment in LD/MD versus HD was transformed by Log_10_. The proteins were organized into families/classes using PANTHER and families/classes were further curated to combine similar PANTHER families and include unassigned proteins to existing families. To find enrichment of EV proteins in infected versus mock samples, the normalized intensity of LD/MD infected proteins was divided by normalized intensity of LD/MD infected proteins and transformed by Log_10_. Heatmaps for enrichment were produced in R using the ComplexHeatmap package and formatted in Adobe Illlustrator.

## Results

### Arabidopsis EV populations can be separated by high-resolution density gradient centrifugation

Purification of EVs from both plant and animal cells commonly involves differential ultracentrifugation followed by floating of EVs in a density gradient (Jeppesen et al., 2019; Kowal et al., 2016; Rutter and Innes, 2017; Temoche-Diaz et al., 2019). First, we wanted to determine if we could enrich for different EV populations simply using differential ultracentrifugation (i.e., by sequential pelleting at increasing speeds). A previous report suggested that EVs pelleted at 40,000 g are enriched in the EV marker protein PENETRATION 1 (PEN1), while EVs pelleted at 100,000 g are enriched in another EV marker protein TETRASPANIN 8 (TET8) (He et al., 2021). To isolate EVs from Arabidopsis, we followed a previously published protocol (Rutter et al., 2017). In brief, 6-week-old Arabidopsis rosettes were vacuum infiltrated with vesicle isolation buffer (VIB). The infiltrated rosettes were gently spun at 700 g to collect the apoplastic wash fluid (AWF) (Figure 1a). After passing the AWF through a 0.22 um filter, the AWF was spun at 10,000 g (P10) in a fixed-angle rotor (TLA100 series) to pellet large particles. The supernatant was spun at either 40,000 g for one hour (P40) or 100,000 g for one hour (P100) using the same rotor. The supernatant of the P40 was moved to a new tube by decanting and was spun at 100,000 g (P100-40) to pellet any remaining EVs left after the P40 step. The resuspended pellets were examined using transmission electron microscopy (TEM), nanoparticle tracking analysis (NTA), and immunoblot. As has been previously observed, 40,000 g was sufficient to pellet EVs from Arabidopsis AWF (Rutter and Innes, 2017; Zand Karimi et al., 2022). TEM, NTA, and immunoblot for the previously described EV markers PEN1, TET8, PATELLIN 1 (PATL1), RPM1-INTERACTING PROTEIN 4 (RIN4), and PENETRATION 3 (PEN3) all indicated the presence of EVs in the P40. The P100-40 contained some residual EVs, but through TEM we found that the P100-40 was mostly enriched in nanofilaments. The P100-40 was also enriched in the proteins ETHIOSPECIFIER MODIFIER 1 (ESM1), RUBISCO LARGE SUBUNIT (RBCL), ACTIN, and ANNEXIN 1 (ANN1) raising the question whether these proteins were genuine EV markers. TET8 seemed to pellet in the P40 just as efficiently as other EV markers, which though contrary to a previous publication, is likely attributable to small discrepancies between protocols, such as the centrifuge tube size, type of rotor used (fixed angle versus swinging bucket) and/or the method employed to remove the supernatant (He et al., 2021). As the TET8 native antibody strongly cross-reacted with a non-specific band at 25 kDa and 55 kDa, we also confirmed this result using a transgenic Arabidopsis line expressing both RFP-tagged PEN1 and GFP-tagged TET8 (Figure 1, Figure S1).

To test whether TET8 and PEN1 mark different populations, as previously reported, we observed the P40 EV pellet from Arabidopsis expressing the TET8-GFP and RFP-PEN1 transgenes using total internal reflection microscopy (TIRF-M) (Figure 1b). Small, bright, particles were observed in EVs isolated from transgenic Arabidopsis plants. We did not observe similar fluorescent particles in concentrated samples of P40 EVs isolated from Col-0 not expressing the transgenes. Co-localization of TET8-GFP and RFP-PEN1 puncta was only observed in 8% of particles (Figure 1c), suggesting that TET8 and PEN1 are generally secreted on separate populations of EVs.

Since the P100 was enriched in all EV markers and contained both vesicles and nanofilaments, we proceeded to separate the diversity of particles and proteins found in the P100 using a high-resolution density gradient. An EV pellet (P100) collected from 100 Arabidopsis rosettes was bottom-loaded beneath a discontinuous iodixanol gradient identical to gradients previously used to separate EV populations collected from human cell lines (Figures 2a and b) (Jeppesen et al., 2019). TEM, NTA, and immunoblots showed that three distinct populations of particles floated at different densities in the gradient: low density (∼1.05-1.08 g/mL), medium density (∼1.10-1.13 g/mL), and high density (∼1.15-1.18 g/mL). The low-density (LD) population contained the putative EV marker proteins PATL1 and PEN1. The medium-density (MD) population also contained PATL1 and PEN1 but was strikingly enriched for the EV marker protein TET8. The high-density (HD) population contained all putative EV markers except TET8, as well as ESM1 and RBCL, which were absent from the LD and MD fractions. The HD fraction was enriched for PEN3, but PEN3 also occurred in LD and MD populations in lesser amounts. TEM showed that the HD fraction was enriched in nanofilaments and other particles <50 nm in diameter.

To conclusively separate LD and MD EVs, we repeated the experiment using a 24-part density gradient (Figure 2c). The 24-part density gradient clearly illustrates the specificity of TET8+ EVs for the MD population in contrast to PATL1+ EVs which are found across all densities. We also observed that though PEN1 appears across a wide range of densities, PEN1 is enriched in the MD population, thus bearing some similarity to TET8+ EVs.

Using a transgenic plant expressing both RFP-PEN1 and TET8-GFP allowed us to probe for more potential EV marker proteins (Figure 2d). Tagged versions of PEN1 and TET8 floated in the gradient as before. ANN1 was present in all three populations, very similar to PEN1 and PATL1. RIN4 was specific to LD and MD populations and did not occur in HD fractions. In this density gradient, we also found that MD EVs had a smaller diameter than other populations of EVs, suggesting that TET8+ EVs may be smaller than other types of EVs. Overall, TET8 was found to be a specific marker of MD EVs and RIN4 was found to be a marker of LD and MD EVs. Other EV marker proteins (PATL1, PEN1, and ANN1) were found across all densities, even high densities which did not exclude other small particles, nanofilaments, and the proteins ESM1 and RBCL.

### Arabidopsis EVs co-pellet with non-EV proteins and cell wall nanofilaments

We hypothesized that ESM1 and RBCL may not be associated with EVs, since they floated exclusively at high densities. We therefore performed a protease protection assay to test whether ESM1 or RBCL are protected from protease digestion by a lipid bilayer (Figure 3a). ESM1 and RBCL1 were digested after 2 hours of incubation with trypsin and chymotrypsin, regardless of detergent treatment (Figure 3b), indicating that these proteins are located outside of EVs. By contrast, most other proteins (TET8, PEN1, RIN4, PATL1, PEN3, and ANN1) were protected from protease in the absence of detergent, suggesting that these proteins are secreted in EVs and thus are good EV marker proteins.

In stain-free gels, we observed a protein band at 42 kDa that was especially abundant in the fraction contain nanofilaments (Figure 2d). We speculated that this band might correspond to ACTIN, which is 42 kD. Previous publications have shown that ACTIN co-purifies with plant EVs (Rutter and Innes, 2017), and that PEN1 is recruited to ACTIN patches at the plasma membrane during infection by powdery mildew (Qin et al., 2021; Yang et al., 2014). We therefore prepared a P100-40 pellet, which is enriched in nanofilaments (Figure 1a), and separated proteins by SDS-PAGE, stained with Coomassie, and excised abundant gel bands of this size. Mass spectrometry identified several ACTIN proteins in these abundant bands (Figure 4a). We next tested whether ACTIN would float with high density fractions that contained nanofilaments. Contrary to our hypothesis, ACTIN was specifically identified only in medium densities (Figure 4b). In this density gradient, we isolated EVs using VIB that contained 10 mM EGTA because Ca2+ can affect actin filament stability (Yin et al., 1981). This change in buffer components had the unintended side-effect of shifting HD EVs to slightly lower densities, suggesting that calcium either in the apoplast or in VIB may cause some EVs to aggregate and float at higher than usual densities.

To directly test whether nanofilaments that co-pellet with EVs were composed of ACTIN, we stained an EV pellet with rhodamine-phalloidin and observed the staining pattern using TIRF-M. Instead of linear filaments, we observed large circular, ring-like structures with the same approximate diameter of EVs (Figure 4c). This result indicates that the nanofilaments are not composed of ACTIN, and together with the previous result showing that ACTIN co-fractionates with MD EVs, instead suggests that ACTIN may bind to the surface of MD EVs.

Since ACTIN did not co-fractionate with nanofilaments in the density gradient, we instead hypothesized that the nanofilaments were composed of cellulose or other cell wall components sloughed from the cell wall during AWF isolation. To test this hypothesis, we digested a P100 (which contains EVs and nanofilaments) with Onozuka R-10, a mixture of cell wall digesting enzymes typically used for protoplast isolation. After 2 hours, no nanofilaments could be observed using TEM (Figure 4d). Along with cellulose, pectin is a major component of the cell wall (Delmer et al., 2024). Therefore, we used the pectin antibody JIM7 to probe for the presence of pectin in EV pellets that contained nanofilaments. Dot blotting showed that pectin is present in roughly equivalent amounts in the P40 and P100-P40, present in the AWF and decreased in the S100, indicating that the majority of pectin in AWF is pelleted at 100,000 g (Figure 4e). Pectin in an EV pellet could be digested with either Onozuka R-10 or pectolyase, confirming the specificity of the JIM7 antibody for pectin (Figure 4e). To assess whether pectin co-purified with filaments in a high-resolution density gradient, we analyzed a density gradient using the JIM7 antibody. This analysis confirmed that pectin is found only in the high-density fractions, which also contain filaments (Figure 4e). We therefore concluded that these nanofilaments are likely derived from the cell wall and are composed at least partially of pectin. Importantly, they can be mostly separated from EVs using a high-resolution density gradient as they equilibrate at higher densities than most EV populations.

### Arabidopsis EV populations respond differentially to hormone treatment and temperature stress

Treatment with the immune signaling hormone salicylic acid (SA) was previously shown to induce secretion of EVs (Rutter and Innes, 2017). To assess whether this was a general EV response, or was restricted to a subset of EVs, we sprayed six-week old Arabidopsis plants with 2 mM SA, 100 µM methyl-jasmonic acid (me-JA), or a mock solution, and then collected P100 pellets at various times after hormone spray. PEN1 and TET8 accumulation in the P100 differed dramatically between hormone treatments (Figure 5a). PEN1 showed a two-fold increase in response to SA at 24 hps as previously described (Rutter, 2017), and a smaller but significant increase in response to me-JA 24 hps. TET8 changed most drastically, showing a striking reduction in response to SA 24 hps and five-fold enrichment in response to JA 24 hps (Figure 5a). TET8 was the only protein to experience a reduction in secretion in response to either hormone. In contrast to TET8 and PEN1, PATL1 had only mild to no response to elicitation. The level of EV proteins in the cell lysate remained constant during elicitation, suggesting that the loading of EV proteins into vesicles must be tightly regulated. We also found that particle concentration, except for 24hr SA treatment, was not significantly increased by SA and JA relative to mock (Figure 5a). Together, this supports a model in which differential loading of EVs occurs during immune hormone signaling.

**FIGURE 5.**
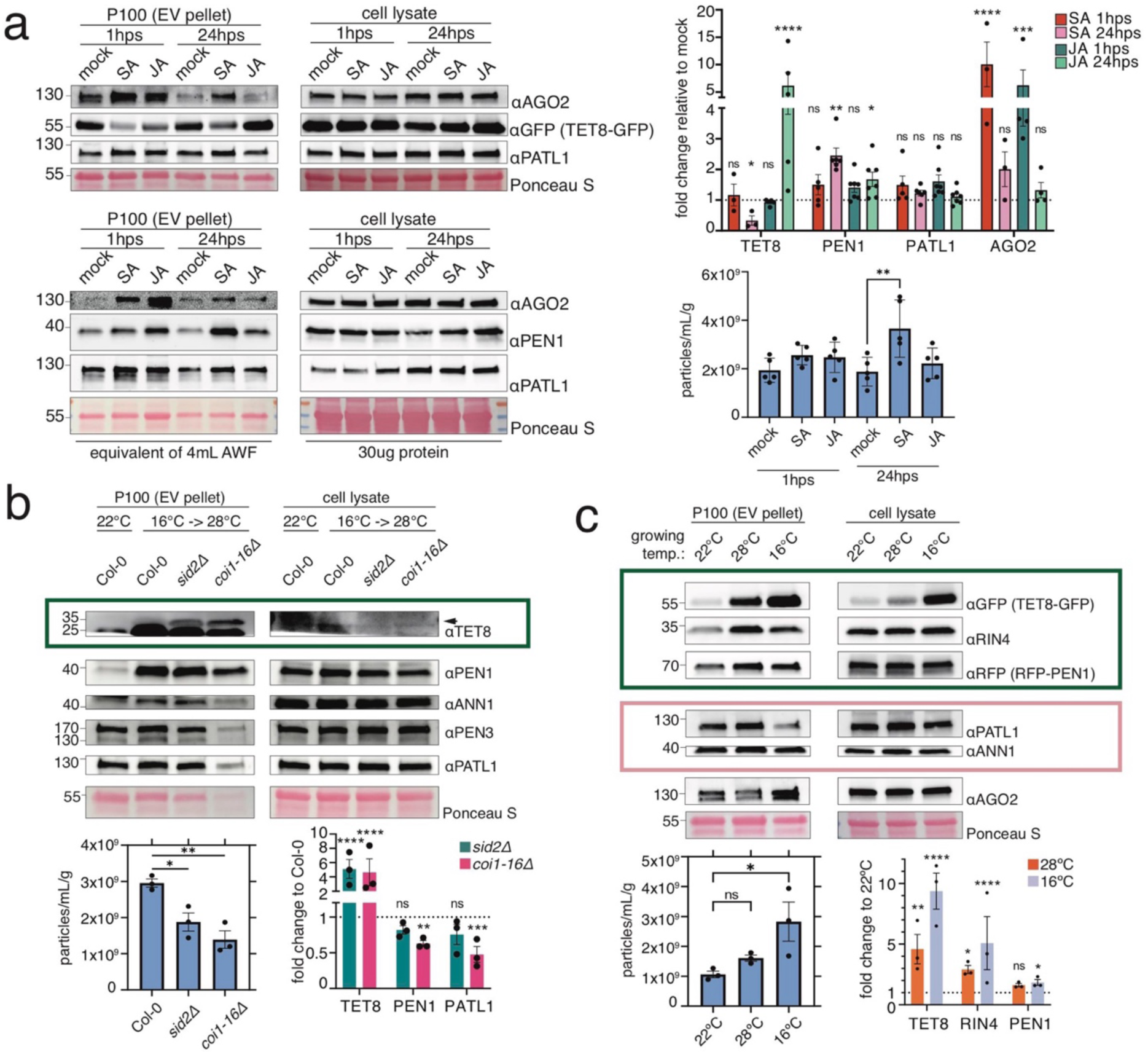
EV population secretion is differentially influenced by immune hormones and temperature. (a) Representative immunoblots, densitometry of immunoblots, and NTA of EV and cell lysate samples collected 1 hour post foliar spray (hps) and 24 hps with 2 mM salicylic acid (SA), 100 µM methyl-jasmonic acid (JA), or buffer without hormone (0.01% Silwet, 20 mM MES, pH 6). For EV samples, the P100 collected from 5 mL of AWF was loaded for each lane. For cell lysate, 30 µg of protein was loaded in each lane. For densitometry, the graph shows the average fold change of each protein relative to that protein’s level at the corresponding mock treatment at that timepoint. Individual points are densitometry values from 3 to 7 immunoblots from independent biological replicates. A two-way ANOVA followed by a post-hoc Dunnett multiple comparison test was used to detect statistical significance between mock protein level and protein level in response to hormone treatment at each timepoint. For NTA, average particle concentration (particles/mL) was determined from 6 independent biological replicates and was normalized using the fresh mass of the plants from which the EV pellet was isolated. A one-way ANOVA was used to detect statistical significance, followed by a post-hoc Sidak multiple comparison between mock and hormone spray at their respective timepoints. (b) Representative immunoblots, densitometry, and NTA of EV and cell lysate samples from indicated genotypes. Plants were grown at either 22°C, or at 16°C and moved to 28°C 3 days before the experiment to disable the *COI1* protein in the *coi1-16* mutant. For EV samples, the P100 collected from 5 mL of AWF was loaded for each lane. The dark green box highlights the unique response of TET8, and the arrowhead indicates the TET8-specific band. For densitometry, the graph shows the average fold change of each protein relative to wild-type Col-0 that experienced the temperature change. Individual points are densitometry values from 3 independent biological replicates. A two-way ANOVA was used to detect statistical significance, followed by a post-hoc Dunnett’s multiple comparison. For NTA, the average particle concentration (particles/mL) was determined from 3 independent biological replicates and normalized using the fresh mass of the plants from which the EV pellet was isolated. A one-way ANOVA was used to detect statistical significance, followed by a post-hoc Dunnett’s multiple comparison. (c) Representative immunoblots, densitometry, and NTA of EV and cell lysate samples from plants grown at 22°C and then shifted to 28°C or 16°C for 3 days. For EV samples, the P100 collected from 3 mL of AWF was loaded for each lane. The dark green box indicates EV proteins whose secretion is strongly stimulated by temperature change, and the light red box indicates non-responsive EV proteins. For densitometry, the graph shows the average fold change of each protein relative to that protein’s level at 22°C. Individual points are densitometry values from 3 independent biological replicates. A two-way ANOVA was used to detect statistical significance, followed by a post-hoc Tukey multiple comparison. For NTA, the average particle concentration (particles/mL) was determined from 3 independent biological replicates and normalized using the fresh mass of the plants from which the EV pellet was isolated. A one-way ANOVA was used to detect statistical significance, followed by a Dunnett’s multiple comparison. P-values: *<0.05, **<0.01, ***<0.001, ****<0.0001, ns=not significant. Error bars indicate ± SEM.

We also used Arabidopsis mutants to test whether these hormones were responsible for governing EV secretion. *SID2* is required for SA biosynthesis and *coi1-16* is a temperature-sensitive mutant of *COI1,* which is required for JA perception (Ellis and Turner, 2002; Sheard et al., 2010; Wildermuth et al., 2001; Xie et al., 1998). The *coi1-16* mutant displayed a marked reduction in most EV proteins in the P100 pellet and reduced EV number when shifted to a non-permissive temperature (28°C; Figure 5b). Notably, TET8 did not follow this pattern, and secretion of TET8 the *coi1-16* mutant was about 5-fold higher than in wild-type Col-0 at 28°C. Mutation of *SID2* had a milder effect on EV number and EV proteins, but TET8 was also elevated about 5-fold in *sid2* plants compared to wild-type Col-0 (Figure 5b). Together, the foliar spray and mutant study suggest that secretion of EV populations are differentially regulated by SA and JA.

We noticed in the above experiments, which required a temperature shift to inactivate COI1, that EV secretion in wild-type Col-0 plants appeared to be stimulated by increased growing temperature (Figure 5b). We followed up on this result by more thoroughly testing which EV proteins were secreted in response to changes in growing temperature. Plants were moved from their typical 22°C growth chamber to 28°C and 16°C growth chambers for 72 hours. NTA results showed that both heat and cold stimulated an increase in EV number, which was much more pronounced in response to cold stress (Figure 5c). The response of TET8 to temperature was the most extreme of the EV marker proteins, with a large increase in TET8+ EV secretion in response to both temperature treatments, particularly cold (Figure 5c), but this increase was also observed in cell lysate, indicating that temperature stress stimulates total TET8 protein accumulation. RIN4+ EVs were also particularly responsive to temperature stress, and PEN1+ EVs also had notable increases in secretion in response to temperature change. The changes in RIN4 and PEN1 secretion occurred in the absence of changes in cellular RIN4 and PEN1 protein levels.

### AGO2 is not secreted in Arabidopsis EVs

We have previously shown that the RNA-binding protein ARGONAUTE2 (AGO2) is secreted and co-pellets with EVs, but is not protected from protease digestion (Zand Karimi et al., 2021). We confirmed that AGO2 is present in the P40 and is not protected from protease digestion (Figures 6a and b). Using an HA-AGO2 Arabidopsis line, we found that AGO2 floats at high densities (Figure 6c), which together with the protease protection assay suggests that AGO2 is not secreted in EVs. Treatment with the immune signaling hormones salicylic acid and jasmonic acid caused increased secretion of AGO2 1 hour after spray (Figure 5a). However, AGO2 does not appear to be enriched in EV pellets relative to total cell lysate, whereas the EV markers PEN1 and especially TET8 are highly enriched in EVs (Figure 6d). Together, this suggests a model in which AGO2 secretion in non-EV particles is strongly and quickly stimulated in response to immune signaling hormones.

**FIGURE 6.**
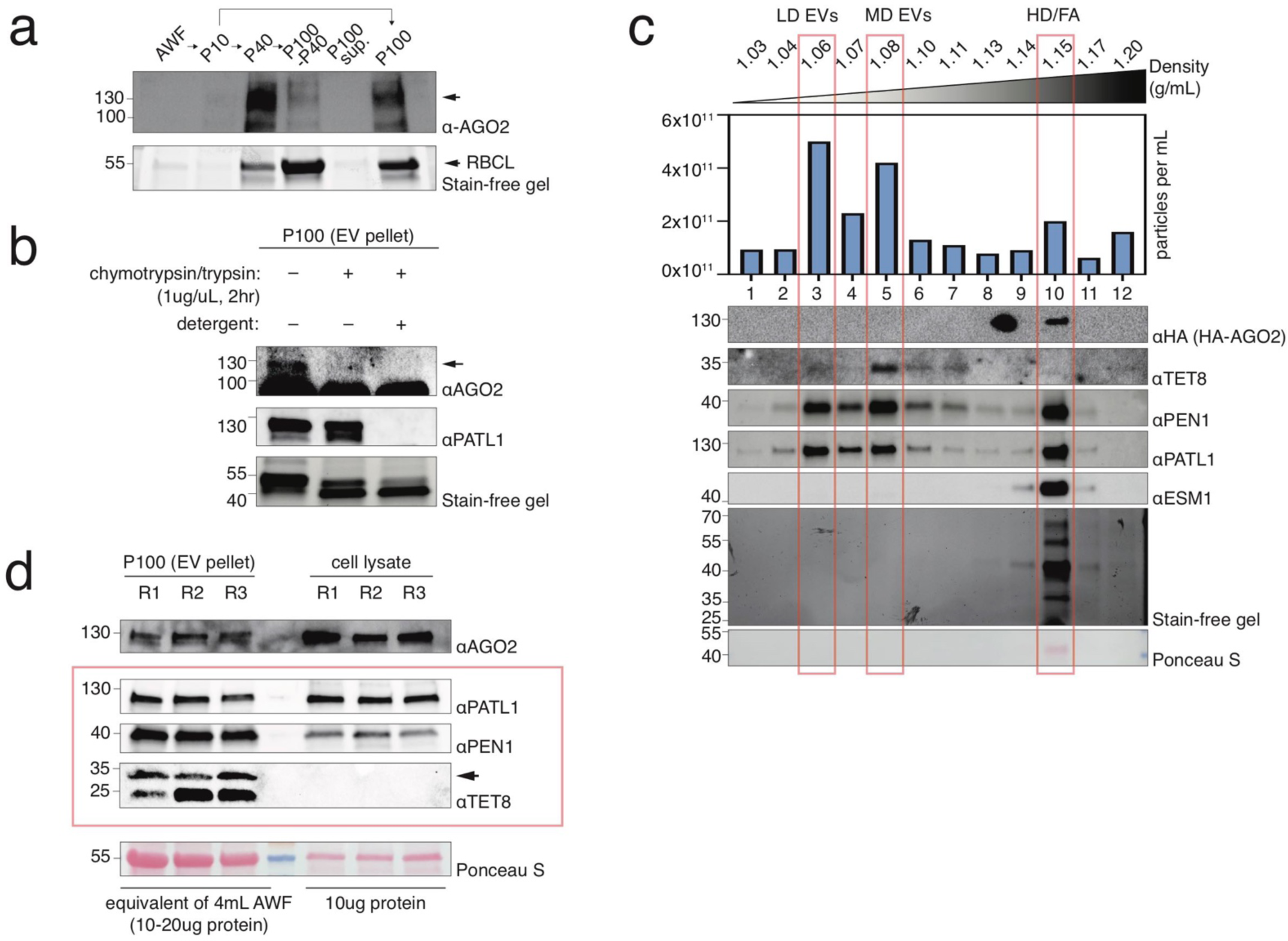
AGO2 co-pellets with EVs but is likely not an EV cargo protein. (a) Differential ultracentrifugation of AWF. EV pellets were each collected from the equivalent of 5 mL of AWF. Arrowhead indicates the AGO2-specific band as verified by probing an *ago2* null mutant (see Figure S1). (b) Immunoblot for AGO2 following protease protection assay. Arrowhead indicates the AGO2-specific band. (c) Immunoblot and NTA following high-resolution density gradient separation of a P100 EV pellet collected from Arabidopsis expressing HA-AGO2 transgene. (d) Immunoblot of AGO proteins and EV marker proteins. For EV samples, the P100 from an equivalent amount of AWF was loaded for each lane, which was between 10-20 µg protein. For cell lysate, 10 µg of protein was loaded in each lane. Arrow indicates the specific bands for AGO2 and TET8 immunoblots. R1, R2, and R3 are replicate 1, 2, and 3.

### Arabidopsis EV populations respond differentially to infection with biotrophic fungi

Previous studies of Arabidopsis EVs have strongly suggested a role for EVs in plant immunity, particularly in defense against fungal pathogens (H et al., 2022; Rutter and Innes, 2018; Rybak and Robatzek, 2019). Therefore, we tested which EV populations responded to the hemibiotrophic fungal pathogens *Colletotrichum higginsianum (Ch)* and *Colletotrichum destructivum (Cd)*, as well as the biotrophic powdery mildew fungus *Golovinomyces cichoracearum (Gc).* Infection with either the compatible *Ch* or the incompatible *Cd*, which is not able to penetrate Arabidopsis, produced an EV response as soon as 16 hours after inoculation (hpi) (Figure 7a). Trypan blue staining of infected leaves showed that 16 hpi with *Ch* was concurrent with spore germination and appressoria formation, while formation of biotrophic hyphae was observed at 64 hpi, as previously described (Figure 7a) (Narusaka et al., 2004). While *Cd* spores germinated and formed appressoria, infection did not progress beyond this point, as previously described (Figure 7a) (O’Connell et al., 2004). The response of TET8+ EVs was strongest, followed by RIN4+ and PEN1+ EVs. Expression of TET8 also seemed to increase in the cell lysate. On the other hand, PATL1, ANN1, and AGO2 did not respond to infection with *Colletotrichum* species.

**FIGURE 7.**
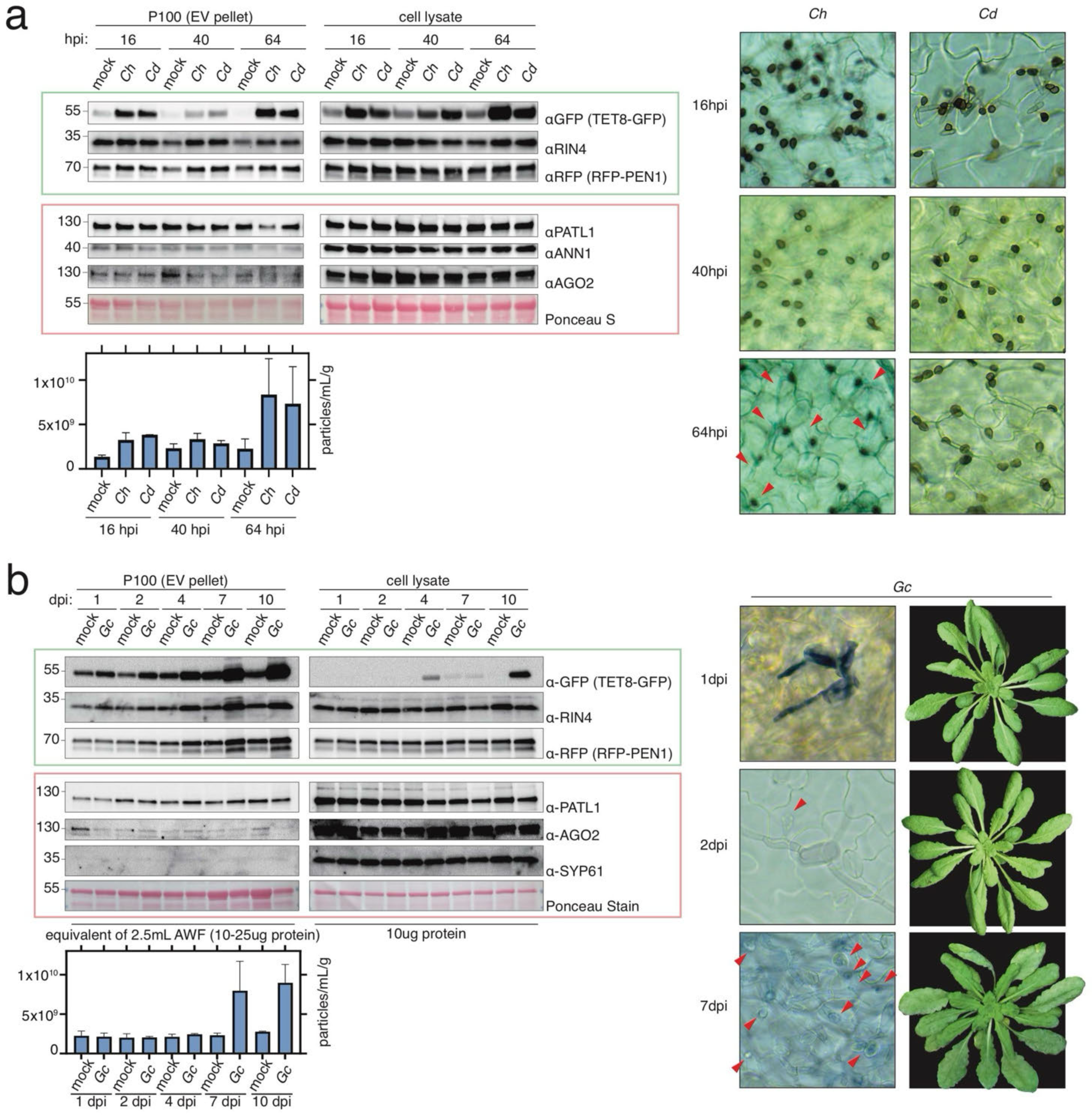
Secretion of EV subpopulations is influenced by infection with biotrophic fungi. Immunoblot, NTA, representative trypan blue stained leaves, and representative photographs of plants at various times after infection with fungi. For (a), Arabidopsis was infected with the compatible *Colletotrichum higginsianum* (*Ch*), the incompatible *Colletotrichum destrucitivum* (*Cd),* or a mock spray. Infection was done by spraying 30 mL of 1x10^7^ spores/mL in distilled water on 72 GFP-TET8 x RFP-PEN1 Arabidopsis plants. Red arrowheads in trypan blue micrograph highlights the development of biotrophic hyphae from melanized appressoria 64 hpi, occurring in *Ch* but not in *Cd* infection. For (b), Arabidopsis was infected with *Golovinomyces cichoracearum (Gc)*. Macroscopic conidia (white powder) became visible 7 dpi. Spores germinated at 1 dpi, formed initial feeding structures at 2 dpi (indicated by red arrowheads), and developed extensive mycelia and large feeding structures 7 dpi. The P100 from an equivalent amount of AWF was loaded for each lane. For cell lysate, an equal amount of protein was loaded per lane. Blots were processed and signal was detected in parallel, so relative enrichment of EV protein compared to cell lysate can be compared. Size of trypan blue micrographs is 110µm x 110µm. Error bars indicate ± SEM. hpi = hours post infection, dpi = days post infection.

Infection with *Gc* produced a similar response, characterized by an increase in secretion of TET8+, RIN4+, and PEN1+ EVs one day after infection which increased throughout infection (Figure 7b). Notably, TET8 was highly enriched in EVs relative to cell lysate, indicating that a large fraction of TET8 is being secreted in response to pathogen infection. Secretion of SYP61, a marker of cytosolic endosomes, was not observed, indicating that cell damage was not likely a source of contamination in EV pellets despite infection. Together, these results suggest that infection with biotrophic fungi induces secretion of a specific population or populations of EVs as soon as 16 to 24 hours after inoculation, which is prior to formation of haustoria, as shown by trypan blue staining of infected leaves. It is also noteworthy that EVs are being isolated primarily from the apoplast, whereas *Gc* infects only epidermal cells. The rapid increase in EV release thus indicates signaling events between epidermal cells and mesophyll cells early in the infection process.

We further investigated the nature of *Ch*’s effect on EV secretion by applying EV pellets collected from mock and infected plants to lower-resolution density gradients and then analyzing two fractions from the gradients using semi-quantitative mass spectrometry (MS). Lower-resolution gradients were used because fractions from high-resolution gradients did not contain enough protein to be analyzed by MS (Figure 8a). We compared proteins found in the low/medium-density (LD/MD) fractions with the high-density (HD) fractions. When we organized the proteins by family/class, we found that protein families associated with membrane trafficking such as small GTPases and SNARE proteins were enriched in LD/MD fractions, while glucosinolate metabolism, RUBISCO, proteasome, and ribosome proteins were enriched in HD fractions (Figure 8b). We also found that LD/MD proteins were about 1.8 times more likely to contain a transmembrane domain than HD proteins, and LD/MD proteins were about 2.6 times less likely to contain a signal peptide than HD proteins. Together, this suggested that our density gradient had successfully purified EVs away from non-EV particles that co-pelleted with EVs. Next, we compared LD/MD proteins from infected and mock plants. Only a few members of each protein family were responsive to *Ch* infection, which included some highlighted EV marker proteins that we had already verified by immunoblot (Figure 8c). These data indicate that secretion of specific EV populations marked by TET8, PEN1, and RIN4, as well as other specific proteins from other families such as RabA family members and EXO70A1, is induced by fungal infection.

**FIGURE 8.**
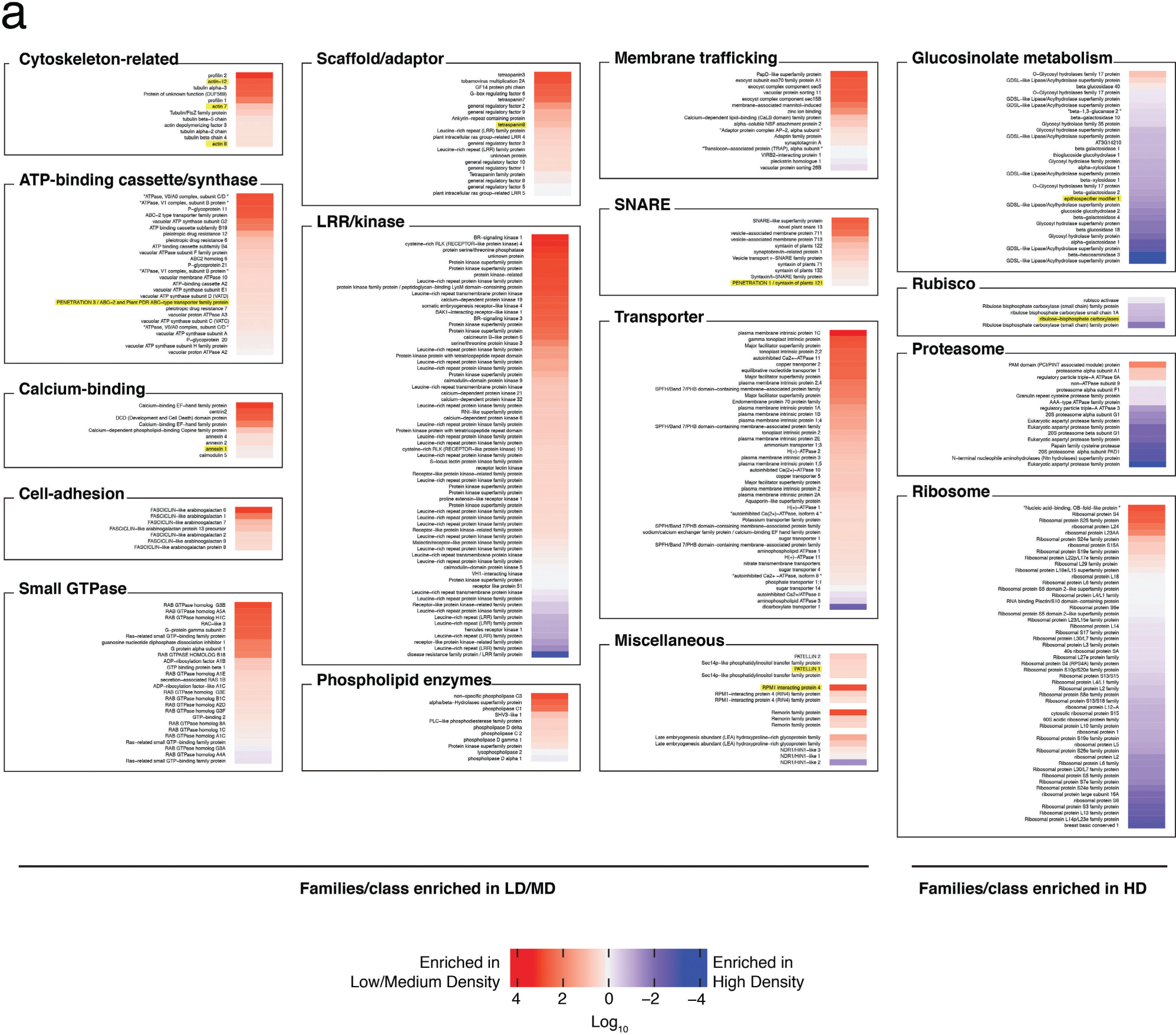
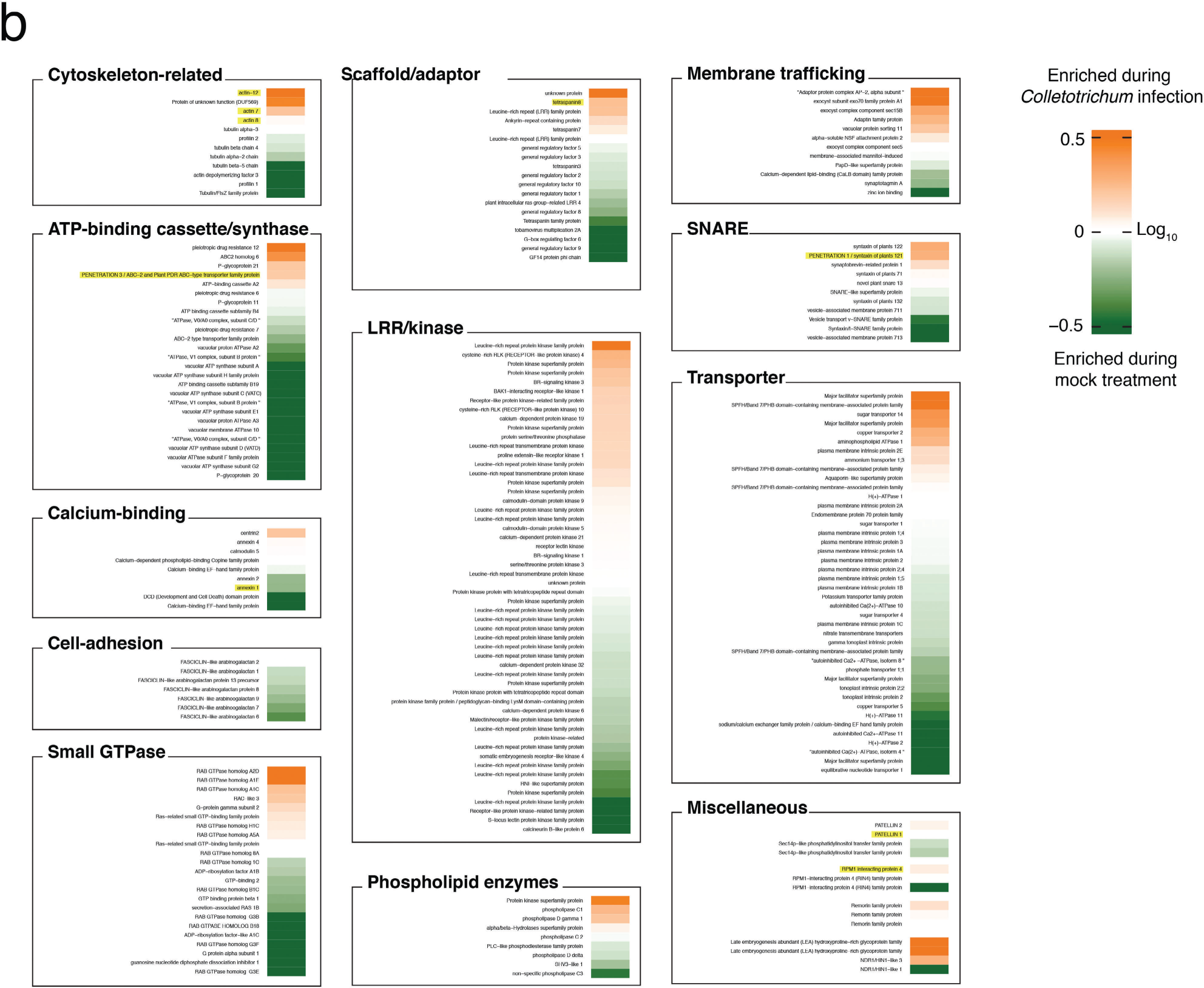
The secretion of a small subset of EV proteins is induced in response to infection. Mass spectrometry (MS) analysis of low-resolution density gradient purified P100 EVs from Arabidopsis plants collected either 64 hours post infection with *Colletotrichum higginsianum (Ch)* or mock. Two independent biological replicates were pooled and analyzed concurrently by MS. Highlighted proteins in (a) and (b) were verified in other independent experiments by immunoblot. In (a), the normalized abundance of proteins in low/medium density (LD/MD) fractions was compared to that of high density (HD) fractions, transformed by Log_10_ such that positive values were enriched in LD/MD and negative values were enriched in HD fractions, and enrichment was plotted as a heatmap. Proteins were organized into families/groups by gene ontology (GO) terms. In (b), the normalized abundance of EV proteins (i.e., proteins enriched in LD/MD fractions) from infected plants was compared to that of mock-treated plants. See Figure S2 for validation of low-resolution density gradient separation of Arabidopsis EVs.

## Discussion

EV populations have been described most extensively in human cell culture. Low-density and high-density subpopulations of EVs have been reproduced in multiple independent studies that utilized iodixanol density gradient purification (Jeppesen et al., 2019; Kowal et al., 2016; Temoche-Diaz et al., 2019). Low-density populations are reported to equilibrate between 1.09 and 1.12 g/mL, while high-density populations equilibrate between 1.14 and 1.16 g/ml (Kowal et al., 2016; Temoche-Diaz et al., 2019). Mass spectrometry and immunoblots showed that low- and high-density EV populations have distinct protein signatures. Low-density EVs are enriched in GO terms for endosome and plasma membrane (Kowal et al., 2016; Temoche-Diaz et al., 2019). Some annexin proteins such as Annexin A1 fractionate at particularly low densities and large diameter EVs marked by Annexin A1 have been observed to bud directly from the plasma membrane using confocal fluorescence microscopy (Jeppesen et al., 2019). On the other hand, high-density EVs are enriched for the GO term endocytic vesicle (Temoche-Diaz et al., 2019). The tetraspanin CD63 was shown to be enriched in high-density EVs and depends on Rab27a for secretion (Temoche-Diaz et al., 2019; Bobrie et al., 2012; Ostrowski et al., 2010). Thus, it has been hypothesized that low-density EVs bud directly from the plasma membrane while high-density EVs are derived from multi-vesicular endosomes (Temoche-Diaz et al., 2019).

Generally, densities from 1.07-1.16 g/mL are associated with the presence of small EVs while densities higher than 1.17 contain non-vesicular particles (Jeppesen et al., 2019; Temoche-Diaz et al., 2019). Non-vesicular particles include ribosomes, proteasomes, extracellular matrix components such as fibronectin, histones, vaults, exomeres, and protein complexes (Jeppesen et al., 2019; Kowal et al., 2016; Zhang et al., 2018). While some cytoskeletal components such as tubulin are non-vesicular, multiple studies have shown that ACTIN is associated with small EV fractions, though immunocapture of tetraspanin+ EVs suggests that ACTIN is not associated with classical exosomes (Jeppesen et al., 2019; Tkach et al., 2017). AGO proteins also float at densities higher than 1.17 and are thus not associated with EVs (Arroyo et al., 2011; Jeppesen et al., 2019; Temoche-Diaz et al., 2019). Up to this point, EV populations in plants have not been well studied. Density gradients using sucrose and iodixanol have been used to purify plant EVs, but density gradients have not yet demonstrated distinct subpopulations of EVs (He et al., 2021; He et al., 2023; Rutter and Innes, 2017). One study reported that differential ultracentrifugation can be used to partially separate PEN1+ and TET8+ EVs (He et al., 2021), and immunocapture of TET8+ EVs has also been used to specifically study the TET8+ EV subpopulation (He et al., 2021; He et al., 2023).

We have found that similar to EVs isolated from human cell culture, bottom-loading Arabidopsis EVs beneath a high-resolution density gradient separates them into distinct populations, which enabled us to define LD EVs (∼1.05-1.08 g/mL), MD EVs (∼1.10-1.13 g/mL), and HD EVs (∼1.15-1.18 g/mL). Like human EVs, LD/MD populations (∼1.05-1.13 g/mL) represent relatively pure EVs as detected by TEM and GO enrichment analysis of LD/MD EV proteomes, while HD EVs (∼1.15-1.18 g/mL) co-fractionated with cell wall nanofilaments, proteasomal and ribosomal components, and AGO2. Like human EVs, MD plant EVs also contain ACTIN. Inclusion of EGTA in the vesicle isolation buffer caused a shift in density of HD EVs and associated particles. Since EVs are negatively charged and cell wall components such as cellulose and pectin are also negatively charged, we speculate that calcium ions may contribute to aggregation of EVs and other particles such as cell wall nanofilaments, changing their apparent buoyant density (Delmer et al., 2024).

Our findings are consistent with the previous report that PEN1+ EVs and TET8+ EVs represent different subpopulations of vesicles (He et al., 2021). We were not able to repeat the partial separation of PEN1+ and TET8+ EVs using differential ultracentrifugation, possibly due to small discrepancies between protocols, such as the centrifuge tube size, type of rotor used (fixed angle versus swinging bucket) and/or the method employed to remove the supernatant. However, high resolution density gradients showed that TET8+ EVs float very specifically with MD EVs while PEN1+ EVs are distributed across a wider range of densities. Additionally, TIRF-M of EV pellets from transgenic Arabidopsis expressing RFP-PEN1 and TET8-GFP showed very little co-localization of PEN1 and TET8. It is possible that this small degree of co-localization can be explained by EVs clumping together, a phenomenon which is apparent when observing EVs with TEM. On the other hand, a small fraction of EVs may genuinely contain both TET8 and PEN1. In human EVs, sorting of proteins into EVs has been proposed to occur somewhat stochastically (Fordjour et al., 2022). Indeed, the tetraspanins CD9, CD63, and CD81, all of which are used to define ‘exosomes’ only partially co-localize, implying that the mechanism by which proteins are sorted into EVs is imperfect or leaky (Han et al., 2021). PEN1 is an abundant protein intrinsic to the plasma membrane, and since the membrane of EVs is likely derived from the plasma membrane, PEN1 may be present at a low frequency among most EV populations (Meyer et al., 2009b). Similarly, PATL1 binds to phosphoinositides that are present in plant EV lipids and in plant plasma membranes, suggesting that PATL1 may also be present in a wide variety of EVs simply because the EV membrane is derived from the plasma membrane regardless of biogenesis pathway (Liu et al., 2020; Peterman et al., 2004; Suzuki et al., 2016). On the other hand, TET8 can bind to glycosylinositolphosphoceramides (GIPCs) which are enriched in the plant EV pool, suggesting a high level of specificity in EV protein and lipid sorting in the case of TET8+ EVs (Liu et al., 2024; Liu et al., 2020). Further experiments are needed to elucidate the mechanism by which proteins are sorted into plant EVs.

Different populations of EVs, either defined by differing densities or by inclusion of a particular EV marker protein, have been hypothesized to originate from different cellular origins (Colombo et al., 2014). We find that lower density EVs are associated with larger diameter (average diameter of fraction 1 = 177 nm), which may indicate that LD EVs are more likely to originate from the plasma membrane as microvesicles, consistent with GO analysis of LD human EVs (Temoche-Diaz et al., 2019). Conversely, MD EVs had the smallest diameter of any fraction (average diameter = 83 nm) and was the only density which included TET8+ EVs. The TET8+ EV population is presumed to represent classical exosomes released from fusion of a multivesicular body with the plasma membrane, due to both the co-localization of TET8 with the Rab5 family GTPase ARA6 which marks late endosomes and the fact that TET8 is the closest plant ortholog of CD63, a marker of classical exosomes in human cells (Boavida et al., 2013; Cai et al., 2018; Jimenez-Jimenez et al., 2019). The flotation of TET8+ EVs specifically at medium densities bears a resemblance to CD63+ EVs (Temoche-Diaz et al., 2019). Neither the TET8+ EV population nor the RIN4+ EV population were found in HD EVs, suggesting that these populations do not readily aggregate or interact with cell wall co-isolates. Other protein markers like the phosphatidylinositol transfer protein PATL1, the annexin ANN1, the ABC transporter PEN3, and the syntaxin PEN1 were found across all densities, and therefore EVs marked by these proteins may originate either from multiple cellular origins, or a single origin capable of producing EVs of a wide range of densities.

Different populations of EVs have been hypothesized to have unique functions. In humans, different populations of EVs as defined by density, protein content, cellular origin, or size have been shown to affect gene expression and/or behavior in recipient cells (Hoshino et al., 2015; Tkach et al., 2017; Willms et al., 2016). In plants, EVs may similarly be involved in signaling, but there is more evidence that plant EVs directly interact with pathogens for defense. PEN1 and PEN3 localize to the papillae beneath sites of attempted fungal penetration of the cell wall (Assaad et al., 2004; Meyer et al., 2009a; Stein et al., 2006; Underwood and Somerville, 2013). Secretion of the TET8+ EVs has been shown to be stimulated by Botrytis infection, and TET8+ EVs can be taken up by Botrytis (Cai et al., 2018; He et al., 2021).

Our results suggest that specific populations of EVs defined by protein markers are a part of the plant response to biotrophic fungal pathogens, including PEN1+, TET8+, and RIN4+ EVs. These subpopulations of EVs fractionate differently in a density gradient, and PEN1 and TET8 do not localize to the same vesicles, suggesting that multiple types of EVs are important during fungal response. These specific EV subpopulations are induced very early in response to biotrophic fungal infection, even before penetration of the cell wall, haustoria formation, and encasement formation, consistent with a role for EVs in aiding papillae formation or other forms of pre-penetration immunity. Secretion of anti-fungal EV subpopulations continues and increases throughout infection, so it is quite possible that EVs also contribute to post-penetration fungal immunity. Alternatively, fungal pathogens may be manipulating the plant into secretion of important defense signaling proteins like RIN4, depleting the cellular pool of RIN4 (Belkhadir et al., 2004; Ray et al., 2019).

EV subpopulations may also be important in response to multiple types of stress since secretion of specific subpopulations was observed during treatment with phytohormones or changes in growth temperature (Table 1). Strong induction of TET8+ EVs in response to jasmonic acid, along with previous studies showing increased secretion of TET8+ EVs during Botrytis infection, suggest that TET8+ EVs are important during necrotrophic fungi infection and responses to wounding (Cai et al., 2018; He et al., 2021). TET8+ EVs have been shown to contain antifungal RNA, and it would be interesting to test whether TET8+ EVs are unique in this regard (He et al., 2021). The capacity for plant EVs to function in systemic signaling, including the acquisition of systemic acquired resistance, particularly considering the specific induction of PEN1+ EVs upon treatment with salicylic acid, should be further investigated (Johansson et al., 2014; Klessig et al., 2018). PATL1 may be useful as a “housekeeping” EV protein, since it is present across a wide range of densities and is generally not reactive to stress.

**Table 1.**
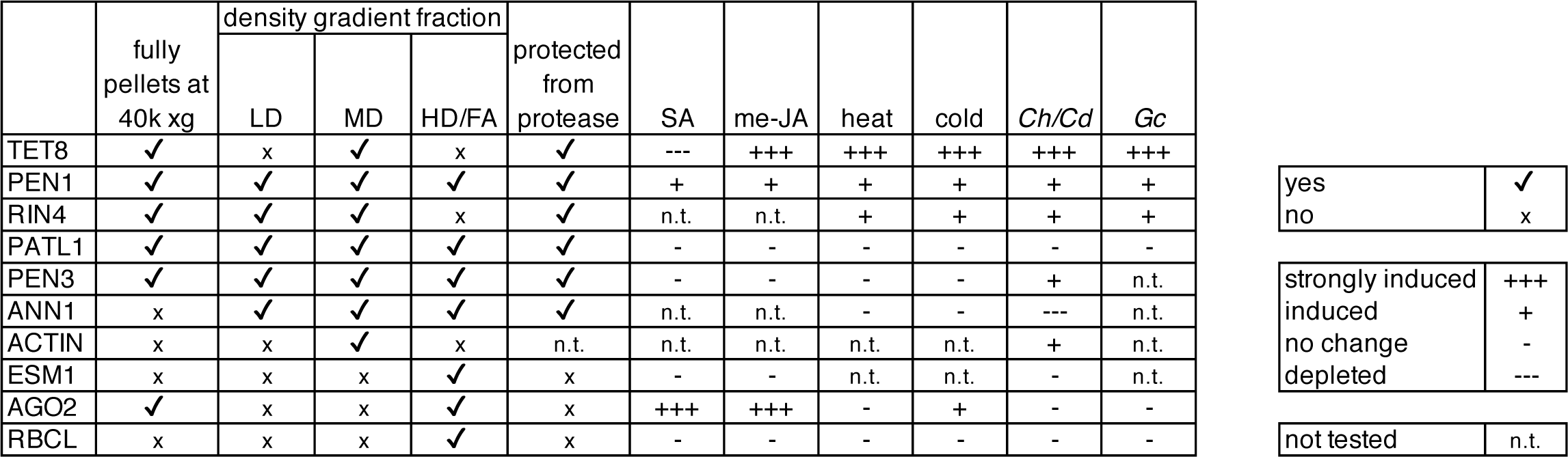
Characteristics of EV marker proteins.

|  | fully<br>pellets at<br>40k xg | density gradient fraction |  |  | protected<br>from<br>protease | SA | me-JA | heat | cold | *Ch/Cd* | *Gc* |
| --- | --- | --- | --- | --- | --- | --- | --- | --- | --- | --- | --- |
|  |  | LD | MD | HD/FA |  |  |  |  |  |  |  |
| TET8 | ✓ | x | ✓ | x | ✓ | --- | +++ | +++ | +++ | +++ | +++ |
| PEN1 | ✓ | ✓ | ✓ | ✓ | ✓ | + | + | + | + | + | + |
| RIN4 | ✓ | ✓ | ✓ | x | ✓ | n.t. | n.t. | + | + | + | + |
| PATL1 | ✓ | ✓ | ✓ | ✓ | ✓ | - | - | - | - | - | - |
| PEN3 | ✓ | ✓ | ✓ | ✓ | ✓ | - | - | - | - | + | n.t. |
| ANN1 | x | ✓ | ✓ | ✓ | ✓ | n.t. | n.t. | - | - | --- | n.t. |
| ACTIN | x | x | ✓ | x | n.t. | n.t. | n.t. | n.t. | n.t. | + | n.t. |
| ESM1 | x | x | x | ✓ | x | - | - | n.t. | n.t. | - | n.t. |
| AGO2 | ✓ | x | x | ✓ | x | +++ | +++ | - | + | - | - |
| RBCL | x | x | x | ✓ | x | - | - | - | - | - | - |

|  |  |
| --- | --- |
| yes | ✓ |
| no | x |
| strongly induced | +++ |
| induced | + |
| no change | - |
| depleted | --- |
| not tested | n.t. |

**Table 2.**
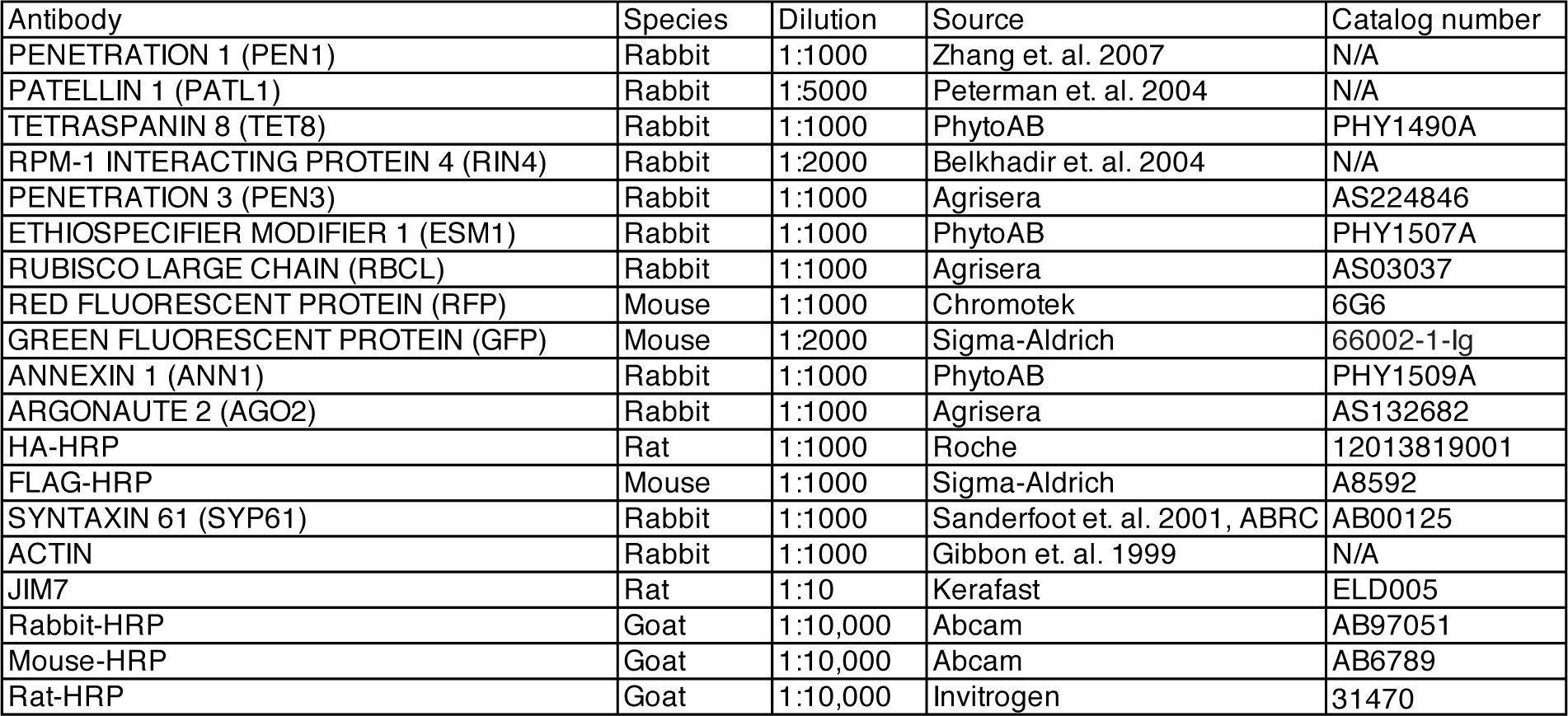
Antibodies used in this study.

The biogenesis of a subclass of anti-fungal EVs may involve actin. Actin patches form at papillae, which are required for PEN3 and PEN1 localization at sites of attempted penetration by powdery mildew (Underwood and Somerville, 2013; Yang et al., 2014). Indeed, actin filament formation is required for immunity to fungal infection (specifically *Cd* infection) (Shimada et al., 2006). Our data indicate that ACTIN is co-secreted with a subpopulation of EVs that are induced by Ch infection, suggesting that actin filaments may be involved in transport or secretion of EVs at sites of fungal infection.

In summary, Arabidopsis EVs are highly diverse and EV subpopulations are reactive to a variety of biotic and abiotic stresses. This work provides a useful collection of protein markers for distinguishing plant EV subpopulations.

## ACKNOWLEDGEMENTS

We thank Mads Nielsen for the TET8-GFP x RFP-PEN1 Arabidopsis line and Janet Braam for the *coi1-16* mutant line, as well as advice for treating plants with methyl jasmonic acid. We also thank the IU Light Microscopy Imaging Center and Andras Kun, the IU Biological Mass Spectrometry Center and Jonathan Trinidad, the IU Physical and Biochemical Instrumentation Facility and Giovanni Gonzalez-Gutierrez, and the IU Electron Microscopy Center and Barry Stein. We are also indebted to Craig Pikaard for access to centrifuges. We also thank Chris Staiger for sharing expertise in TIRF microscopy and providing phalloidin stains and anti-Actin antibody. Thanks also to Shunyuan Xiao for providing powdery mildew strain UCSC1 and to Meenu Singla for validating antibodies using Arabidopsis mutants. B.L.K. was supported by a Carlos O. Miller Graduate Fellowship, the Colonel Bayard Franklin Floyd Memorial Fund, and the William R. Ogg Fellowship, all from the IU Foundation. Additional support was provided by the Cox Research Scholar Program and the IU Undergraduate Research STEM Summer Research Program for funding of undergraduate researchers. This work was supported by grants from the National Science Foundation Plant Biotic Interactions and Plant Genome Research programs (grant numbers IOS-1645745 and IOS-1842685 to R.W.I.) The IU Light Microscopy center’s purchase of the GE Deltavision OMX SR super-resolution microscope was supported by a grant from the National Institute of Health (grant number NIH1S10OD024988-01.)

## AUTHOR CONTRIBUTIONS

Benjamin L. Koch: Conceptualization; Formal analysis; Investigation; Methodology; Visualization; Writing – original draft; Writing– review & editing. Brian D. Rutter: Formal analysis; Investigation; Methodology; Visualization; Writing – original draft; Writing – review & editing. Roger W. Innes: Conceptualization; Funding acquisition; Project administration; Resources; Supervision; Writing – original draft; Writing – review & editing

---

**FIGURE S1.**
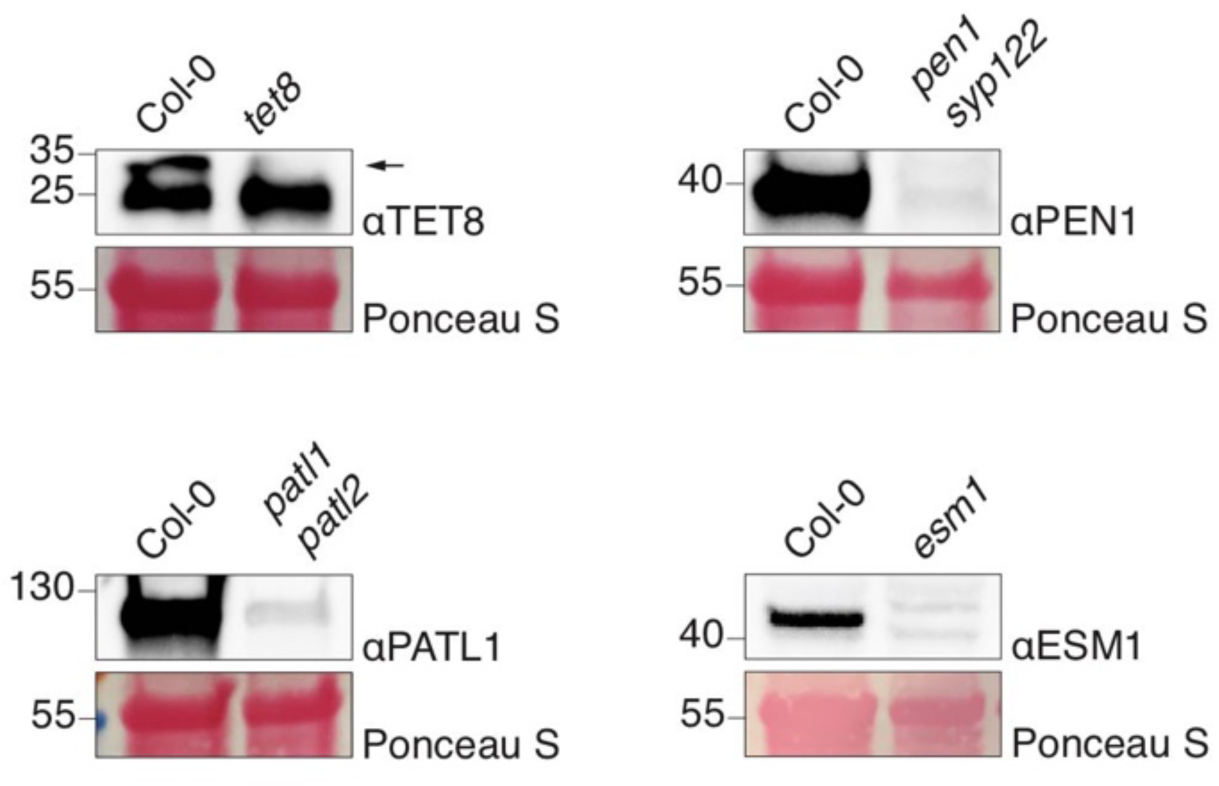
Validation of Arabidopsis antibodies. Equal amounts of cell lysate protein from Col-0 and Arabidopsis mutants were separated by SDS-PAGE and probed with EV antibodies used in this study (see Table 2). The arrow indicates the specific TET8 band.

**FIGURE S2.**
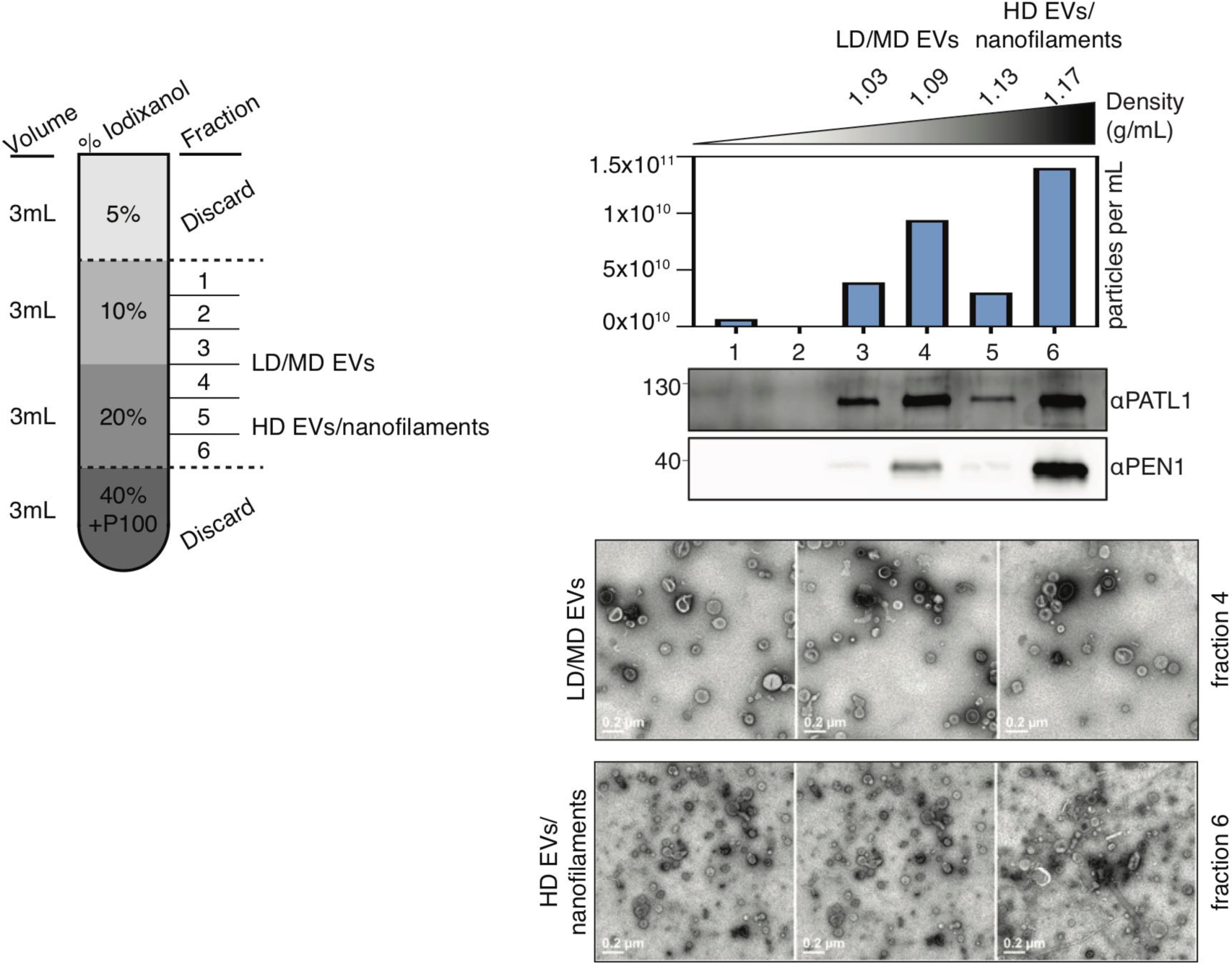
Low-resolution density gradient separation of Arabidopsis EVs for mass spectrometry analysis. NTA, immunoblot, refractometry, and representative TEM images of fractions of a low-resolution density gradient layered above an Arabidopsis EV pellet. From these data, we determined that fractions 3 and 4 contained low-density (LD) and medium-density (MD) EVs, while fractions 5 and 6 contained high density (HD) EVs and nanofilaments.

